# Satb1 integrates cohesin mediated genome organization and transcriptional regulation during T cell development

**DOI:** 10.1101/2025.06.19.657327

**Authors:** Ayush Madhok, Indumathi Patta, Isha Desai, Mohd Tayyab, Vidur Chaudhury, Sushma Vishwakarma, Shagnik Saha, Riitta Lahesmaa, Geeta Narlikar, Sanjeev Galande

## Abstract

Three-dimensional (3D) genome folding, which is highly cell type-specific, plays a crucial role in orchestrating spatiotemporal gene expression. Although factors such as CTCF have been extensively studied in the hierarchical regulation of 3D chromatin organization, the mechanisms driving dynamic genome folding during T cell fate transitions remain incompletely defined. In this study, we reveal that Satb1, a chromatin organizer enriched in the T cell lineage, co-occupies genomic regions with the cohesin complex and Ctcf in double-positive (DP) thymocytes, where chromatin interactions are notably increased. We show that Satb1 physically interacts with the cohesin subunit Smc1a, and its deletion results in aberrant Smc1a binding and reduced chromatin contacts at sites co-occupied by Satb1 and cohesin. In both DP and immature CD4 single-positive (SP) T cells, Satb1 is essential for proper T cell activation and cytokine signaling. At the *Cd3* locus, Satb1 and cohesin collaboratively regulate gene expression, with Satb1 loss leading to disrupted Smc1a occupancy and compromised chromatin interactions. Furthermore, Satb1 shows *in vitro* properties consistent with liquid-liquid phase separation, and disease-associated mutations impair these properties. Together, our findings uncover a molecular mechanism in which Satb1 facilitates chromatin looping through direct interaction with the cohesin complex and its ability to form nuclear condensates, thereby governing transcriptional regulation during T cell development.

## Introduction

Cell fate decisions are primarily governed by cell type–specific epigenetic landscapes and temporally regulated expression of transcription factor (TF) networks. TF binding is constrained by dynamic chromatin states and higher-order chromatin structures [1]. Among these, topologically associating domains (TADs) act as 3D genomic structures that sequester regulatory elements (e.g., enhancers, promoters, non-coding RNAs), thereby providing insulation from neighboring TADs or loop domains [2–4]. An increasing body of evidence highlights the importance of these self-organizing chromatin structures (TADs, sub-TADs) in regulating gene expression and maintaining cellular identity [1, 5, 6]. Proteins like CCCTC-binding factor (CTCF) and the cohesin complex are often enriched at TAD boundaries [7–9] and are implicated in TAD formation via loop extrusion mechanisms [9–11]. While critical, CTCF removal affects only a subset of domain boundaries [12, 13] and has minimal global impact on gene expression [14]. In contrast, depletion of cohesin components abolishes most loop domains but enhances genomic segregation into compartments [14, 15]. Notably, 10–40% of TAD boundaries vary between cell types, contributing to the plasticity of 3D chromatin architecture and lineage-specific transcription programs [16, 17]. T cell development serves as an ideal model to study such variation [18].

In double-positive (DP) thymocytes—where T cell receptors (TCRs) are fully rearranged—the TCR signal induces expression of multiple lineage-determining TFs. Among them, SATB1 (Special AT-rich Binding Protein 1) is an early TCR-responsive TF [19–21] and a T cell–enriched chromatin organizer. SATB1 is critical for thymocyte maturation; SATB1-deficient mice exhibit a developmental block at the DP stage, with markedly reduced CD4^+^ T cell output [22, 23]. Beyond thymic function, SATB1 is also essential for embryogenesis [24], hematopoiesis [25, 26], and regulatory T cell differentiation through its control over the super-enhancer of forkhead box protein P3 (Foxp3) [27]. Owing to its high expression in T cells, SATB1 has been extensively studied for its roles in DP differentiation [27], autoimmunity [28], and T cell migration [29].

SATB1 acts contextually both as an activator and a repressor, depending on post-translational modifications [19, 30], and it interacts with chromatin remodelers (e.g., PCAF/p300, HDACs [31, 32], SWI/SNF [33] and TFs such as β-catenin [34, 35] and CTBP1 [30]. SATB1 has also been linked to CD4/CD8 lineage decisions [27], Th2 differentiation [32, 36], and chromatin looping at loci like MHC Class I [37] and Rag1 [38]. Recent findings reveal its role in sustaining DP identity via regulation of DP-specific super-enhancers [39] and locus-specific chromatin reorganization [40]. However, SATB1’s function during DP fate specification remains incompletely understood.

TFs and other regulatory proteins often aggregate around gene regulatory regions, forming concentrated hubs within the nucleus [41]. Despite CTCF or cohesin depletion disrupting topological insulation, global transcription remains largely unaffected, suggesting that TADs may not be strict barriers to enhancer activity [12, 14, 42]. Nevertheless, clustering of functionally related regions near specific nuclear compartments may enhance transcriptional efficiency by increasing local concentrations of regulatory components [43].

Recent studies suggest that phase separation plays a key role in organizing the 3D genome [41, 44]. Phase-separated condensates—membrane-less, liquid-like assemblies of self-associating proteins—can enrich specific molecular interactions while excluding others [45, 46]. These condensates are now recognized as widespread biological structures implicated in chromatin architecture and transcriptional regulation [46–48].

In this study, we investigated the interplay between Satb1, Ctcf, and cohesin in organizing chromatin interactions in DP thymocytes. We show that Satb1 physically interacts with the cohesin subunit Smc1a. We further examine how Satb1 deletion affects Ctcf/cohesin binding patterns, epigenetic marks, and transcriptional outputs. Additionally, we explore SATB1-mediated transcriptional regulation in both DP and CD4 single-positive (CD4SP) thymocytes. Lastly, we provide evidence that SATB1 can undergo phase separation and identify specific residues essential for condensate formation.

## Results

### Satb1 co-occupies genomic regions with Ctcf and Cohesin in DP thymocytes

To explore the functional significance of Satb1-mediated genome organization during T cell development, we first analyzed publicly available ChIP-seq datasets for Satb1, Ctcf, and cohesin (Smc1a), along with Hi-C data from double positive (DP) thymocytes, where Satb1 expression is markedly elevated (Figure S1A). Using Smc1 HiChIP data from the DP stage [49], we mapped cohesin-mediated chromatin loops to examine the spatial association between Satb1 binding and higher-order chromatin architecture. At the *Ets1* locus, critical for CD8 lineage commitment in the thymus [50], we observed co-occupancy of Satb1 and Ctcf with cohesin loops (Figure 1A). Satb1 was found to associate with a subset of cohesion-mediated loops in DP cells, some independently and others co-bound with Ctcf at one or both loop anchors. A gene-centric analysis based on nearest transcription start site (TSS) revealed that Satb1 and Ctcf share approximately 60% of their bound genes (Figure 1B and S1B). At the *Cd8* super-enhancer (SE) region, Satb1 is found in conjunction with either Ctcf or cohesin (Figure 1C). Based on these genome-wide occupancy profiles we categorized binding sites into Ctcf-only, Satb1-only, and Satb1+Ctcf bound groups and found that around 25% of Satb1-bound regions are shared with Ctcf (Figure S1C). Motif enrichment analysis using MEME Suite [51] revealed that co-bound regions are enriched for both Ctcf and Satb1-like motifs (identified as Onecut), indicating potential direct DNA binding by Satb1 alongside Ctcf (Figure 1D). Similarly, Smc1a sites overlapping with Satb1 also exhibited enrichment for both Ctcf and Satb1-like motifs (Figure S1D), suggesting Ctcf’s involvement in Satb1-cohesin co-occupancy. Using Hi-C data spanning key thymocyte developmental stages, from early thymic precursors (ETP) to mature CD4 cells [52]. We next assessed long-range chromatin interactions involving regions bound by Satb1 alone, Ctcf alone, or both (Figure 1E). While Ctcf-only sites maintained relatively stable interaction frequencies across development, regions co-bound by Satb1 and Ctcf exhibited a marked and progressive increase in long-range interactions during post-commitment differentiation (DN3 onwards). Notably, Satb1-only regions also exhibited a consistent increase in chromatin contacts from the DN3 to DP stages, presumably as a result of increased Satb1 expression in these stages (Figure S1A). Together, these findings suggest that Satb1 association with the chromatin architectural proteins such as Ctcf and cohesin may play a key regulatory role in shaping 3D genome organization during T cell maturation.

**Figure 1.**
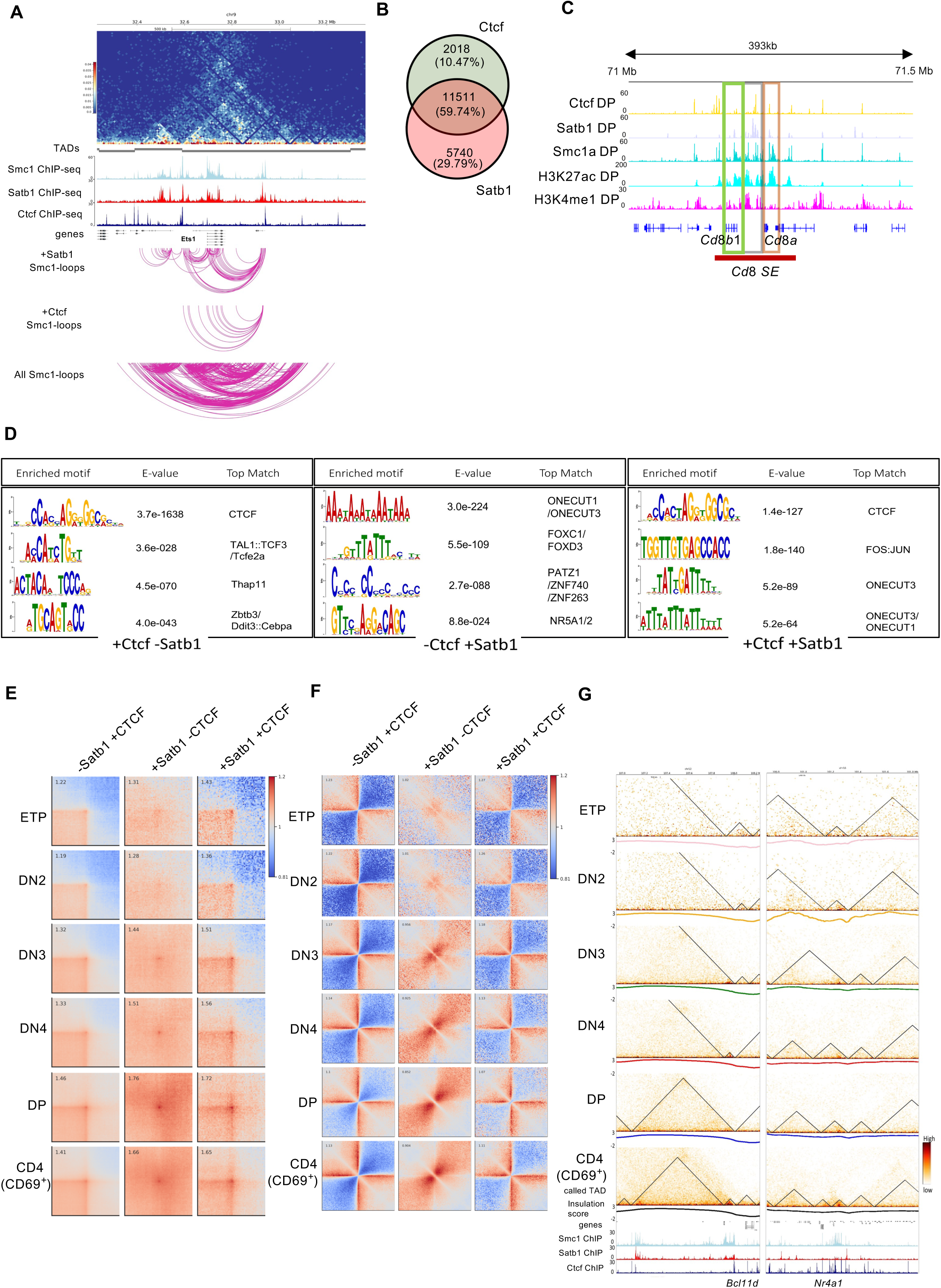
Satb1 co-occupies genomic regions with Ctcf and Cohesin, and binds a subset of insulated regions in DP thymocytes. A. Genomic interactions were depicted at. 1.2 Mb region spanning the *Ets1* locus on Chromosome 9 in DP thymocytes. Top panel: shows HiC contact matrix obtained using publicly available datasets [52]. The called TADs (depicted in grey) show variable bin size dependent on the contact frequency. The genomic region was overlayed with ChIP-seq of Smc1, Satb1 and Ctcf. The following tracks show Smc1 loops co-occupied by Satb1, Smc1 loops bound by Ctcf and total Smc1 loops at the Ets1 locus. B. Venn diagram showing overlap of genes bound by Satb1 and Ctcf in DP thymocytes. Percentages were calculated as ratios to total number of peaks in the gene set. C. Integrative Genomics Viewer (IGV) tracks spanning *Cd8* super enhancer (SE) region showing one dimensional genomic occupancy of Ctcf, Satb1 and Smc1a along with enhancer specific histone modifications. Boxed regions indicate co-bound and exclusively bound regions by these factors. D. Motif enrichment analysis of ChIP-seq peaks; Ctcf excluding Satb1, Satb1 excluding Ctcf, and the shared peaks of the two proteins, respectively (see methods). Top motifs are shown as sequence-logos for each subset along with their statistical probabilities (E-value) and best matched known factors. E. Pileup analysis of developmental stage-specific HiC data. Interactions specific to either Ctcf, Satb1 or both, respectively are averaged over regions from 100 kb to 600 kb. F. Local pileup interactions of HiC data across thymocyte development are depicted as heatmaps. The interactions are calculated for Satb1 alone, Ctcf alone and Satb1-Ctcf co-bound sites. A 600 kb flanking region around different peak sets is depicted as horizontal and vertical lines, marking boundary interactions. G. Snapshot of genomic interactions in various thymic stages at the Bcl11b (left) and Nr4a1 (right) loci. The HiC interactions in different stages are normalized to sequencing depth. Below each HiC contact matrix, the insulation score is plotted which were also used to find TADs (shown as triangles on the contact matrix) using HiC explorer [89]. The following panels show genome browser view of Ctcf, Cohesin (Smc1a) and Satb1 binding events in DP thymocytes. The ChIP signal was normalized by reads per genomic content (RPGC) using mm10 build.

### Satb1 co-occupies TAD boundaries along with Ctcf in thymocytes

We next examined the average local chromatin interactions during T cell development associated with regions bound by Ctcf alone, Satb1 alone, or co-bound by both Satb1 and Ctcf. In early developmental stages (ETP to DN3), regions co-bound by Satb1 and Ctcf exhibited fewer upstream and downstream interactions (Figure 1F). However, at the DN4 and DP stages, these co-bound sites showed a progressive loss of insulation, particularly at Satb1+Ctcf sites, suggesting an increase in local interactions flanking these formerly insulated domains (Figure 1F). In contrast, Ctcf-only regions maintained strong insulation across all stages, with only a modest increase in local interactions observed at the DP stage. Interestingly, Satb1-only regions displayed a marked rise in 3D interactions beginning at the DN3 stage, though these interactions were not enriched at domain boundaries. This suggests that Satb1’s boundary-related activity may depend on co-association with Ctcf (Figure 1F). Furthermore, we calculated boundary strength of insulation boundaries found in DPs using the diamond insulation method [53] and mapped the association of both Satb1 and Ctcf to these boundaries. While Satb1 was present at both strong and weak boundaries, its signal was lower compared to Ctcf (Figure S2A). We next examined the extent to which Satb1 binding sites overlapped with TAD boundaries identified during post-commitment stages of T cell development (Figure S2B). Around 20% of TAD boundaries in DP cells were associated with Satb1, many of which were unique and not shared with Ctcf. Notably, Satb1 co-occupied around 25% of Ctcf-defined boundaries in both DP (Figure S1E) and CD4 (Figure S1F) thymocytes. Furthermore, about 55% of genes located at TAD boundaries in DP cells were also direct Satb1 targets (Figure S2C). Genome-wide analyses revealed that Satb1 is more strongly enriched at DP-specific boundaries compared to adjacent genomic regions (Figure S1G). Finally, by integrating stage-specific insulation profiles with Satb1, Smc1a, and Ctcf binding data, we observed dynamic changes in boundary strength at key T cell regulatory genes. From the DN4 to DP transition, several boundaries exhibited altered interactions between neighboring domains (Figure 1G and Figure S2D-F), coinciding with increased gene expression and strong Satb1 binding. These findings suggest that Satb1 may facilitate stage-specific chromatin remodeling and fine-tuned regulatory interactions essential for T cell development.

### Satb1 physically interacts with cohesin subunit Smc1a in DP thymocytes

Satb1, known for its specific affinity to base-unpairing AT-rich sequences [54, 55], recruits a range of transcription factors [35, 37] and chromatin-modifying enzymes [19, 30, 32, 33] to its genomic targets. Both earlier and more recent studies have suggested that multiple regulatory proteins act in concert to shape chromatin architecture [1, 56–58]. For example, the mediator and cohesin complexes have been implicated in linking genome topology with transcriptional activation in certain contexts [59]. Given that Satb1 co-localizes with both Smc1a and Ctcf at specific genomic sites (Figure 2A and S3A–B), we sought to determine whether SATB1 physically associates with SMC1a, CTCF or other chromatin-associated factors. To address this, we conducted immunoprecipitation followed by mass spectrometry (IP-MS) using lysates from DP thymocytes. This analysis identified approximately 250 high-confidence Satb1-interacting proteins, with a confidence threshold above 90%. Protein–protein interaction (PPI) network analysis using Cytoscape (p < 0.05) grouped these interactors into functionally enriched clusters (Figure 2B), with prominent categories including chromatin organization/modification, nucleosome assembly, and co-translation. Notably, Smc1a was among the most highly enriched interactors. To validate this interaction in T cells, we performed super-resolution confocal imaging of Satb1 and Smc1a in DP thymocytes (Figure 2C). Satb1 is known to exhibit a distinctive cage-like nuclear distribution in thymocytes [33, 35], and interestingly, Smc1a displayed a similar pattern. Z-stack analysis further revealed that Satb1 and Smc1a co-localize throughout the nucleus in distinct punctate structures (Figure 2D). Co-immunoprecipitation assays demonstrated a physical interaction between SATB1 and SMC1a (Figure 2E–F). We also observed an interaction between Satb1 and Smc3, another core subunit of the cohesin complex (Figure S3C). Interestingly, virtually no interaction was detected between Satb1 and Ctcf in DP thymocytes (Figure S3D). Further evidence of functional cooperation between Satb1 and the cohesin complex comes from transcriptomic analyses: a significant overlap was seen in the set of genes dysregulated in Rad21-deficient (cohesin-deficient) and Satb1-deficient DP thymocytes (Figure S3E). These shared targets were enriched for signaling pathways such as TCR and MAPK (Figure S3F), indicating potential co-regulation by Satb1 and cohesin.

**Figure 2.**
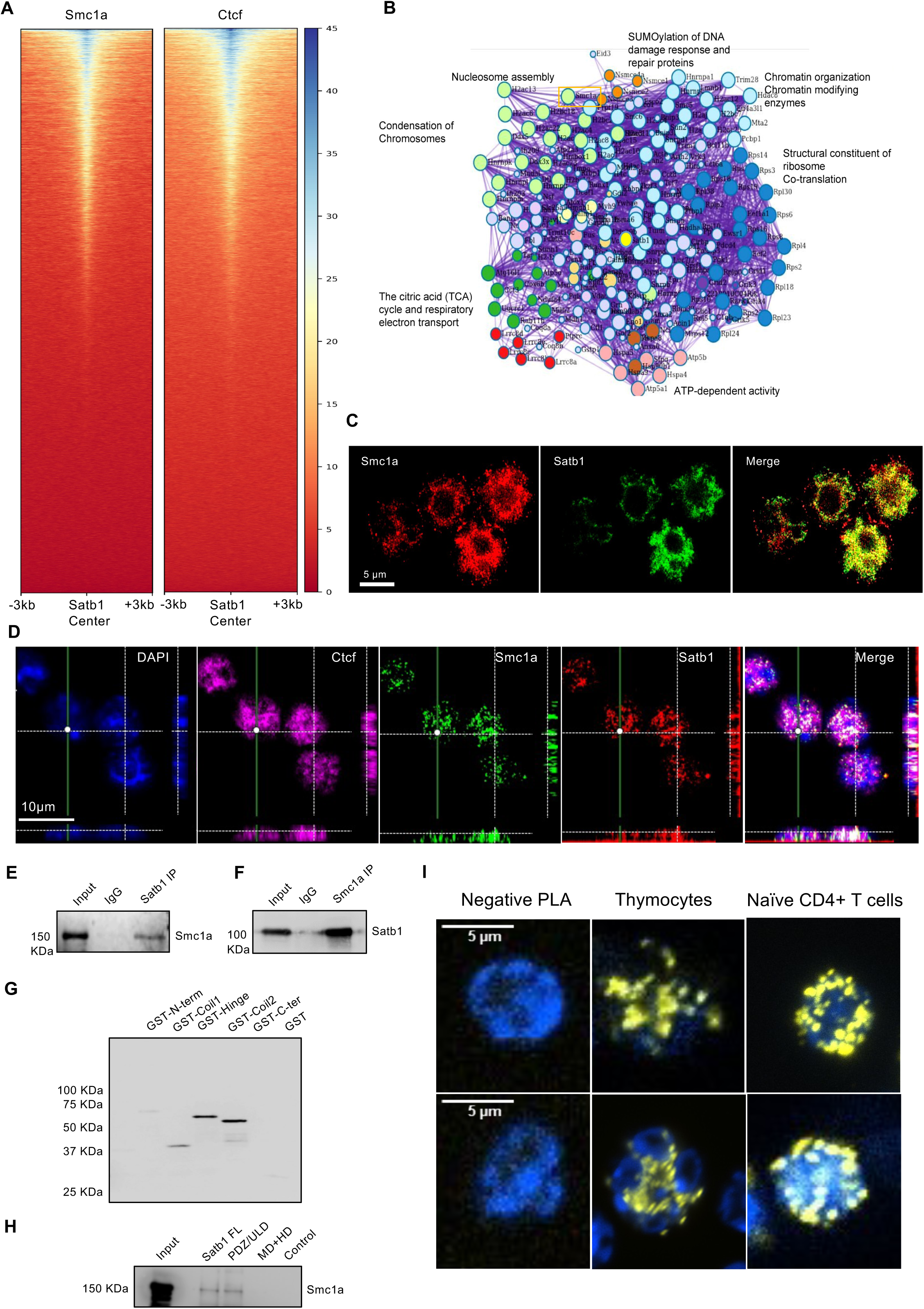
Satb1 directly interacts with Cohesin in DP thymocytes. **A**. Heatmap of normalized ChIP seq signal of Smc1a (left) and Ctcf (right) relative to Satb1 binding sites in DP thymocytes was shown. **B.** Protein lysate from DP thymocytes was prepared. Immunoprecipitation (IP) of Satb1 was carried out and mass-spectrometry (MS) was performed on the IP-ed proteins. Proteins enriched significantly (>90%) with at least 3 peptides were selected for the protein network analysis. The network displayed shows the most significantly enriched processes in which Satb1 interactor proteins are involved. Smc1a (boxed), was detected as a Satb1 interacting protein. **C.** Co-localization of Satb1 and Smc1a was confirmed in DP thymocytes using super-resolution microscopy. Smc1a was labelled with Alexa-fluor 594 (shown in red). Satb1 was labeled with Alexa-fluor 488 (shown in green). The merged image reveals their interaction foci (yellow), which displays the characteristic ‘cage-like’ pattern of Satb1 localization in DP thymocytes [77]. **D.** Immunostaining of Satb1 (red), Smc1a (green), Ctcf (magenta) and DAPI (blue) was carried out in DP thymocytes. z-sectioning shows that Satb1 and Smc1a co-localize throughout the nuclear depth. **E** and **F.** Satb1 interaction was confirmed with Smc1a via co-immunoprecipitation (Co-IP). Satb1 IP was performed, followed by Smc1a western blotting (WB), and vice-versa. **G**. In vitro pull-down assay was carried out using full length HIS-tagged Satb1 with individual domains of Smc1a. Ni-NTA pull down (PD) followed by western blotting of the PD with anti-GST antibody is shown. **H.** Like **G**, a reverse assay was carried out in which full length Smc1a was incubated with the ULD/PDZ and MD+HD domains of Satb1, individually. HIS-tag pull down followed by western blotting of Smc1a is shown. **I.** Confocal images at 100X magnification showing amplification of colocalization signal (yellow) by proximity ligation assay (PLA). DNA was stained with DAPI (blue). Images from two separate experiments are shown as top and bottom panels.

To dissect the molecular interface of the Satb1–Smc1a interaction, we engineered domain-specific constructs of both proteins, tagging Satb1 with polyhistidine and Smc1a with GST (schematically shown in Figure S4A–B). In vitro pull-down assays using full-length and truncated proteins (Fig. 7B, S4C) revealed that Satb1 strongly interacts with the coil-II and C-terminal domains of Smc1a, while showing weaker interaction with the hinge region (Figure 2G). Conversely, Smc1a interacts specifically with the N-terminal ULD/PDZ-like domain of Satb1, which is known as its principal protein–protein interaction domain (Figure 2H). These results suggest that the Satb1–Smc1a interaction primarily involves the N-terminal region of Satb1 and the C-terminal region of Smc1a This is further supported by in silico structural predictions of the Satb1–Smc1a heterodimer (Figure S4D), which are consistent with the data shown in Figure 2G and E. We also performed proximity ligation assay (PLA) to ascertain the extent of physical association between Satb1 and Smc1a (Figure 2I). We observed specific points of high proximity between these two proteins in thymocytes. Interestingly, we also found their interaction also in naïve CD4 T cells (Figure 2I). Collectively, these data indicate that Satb1 shares its genomic occupancy with cohesin through protein-protein interaction in thymocytes and mature T cells, enabled through its ULD domain.

**Figure 3.**
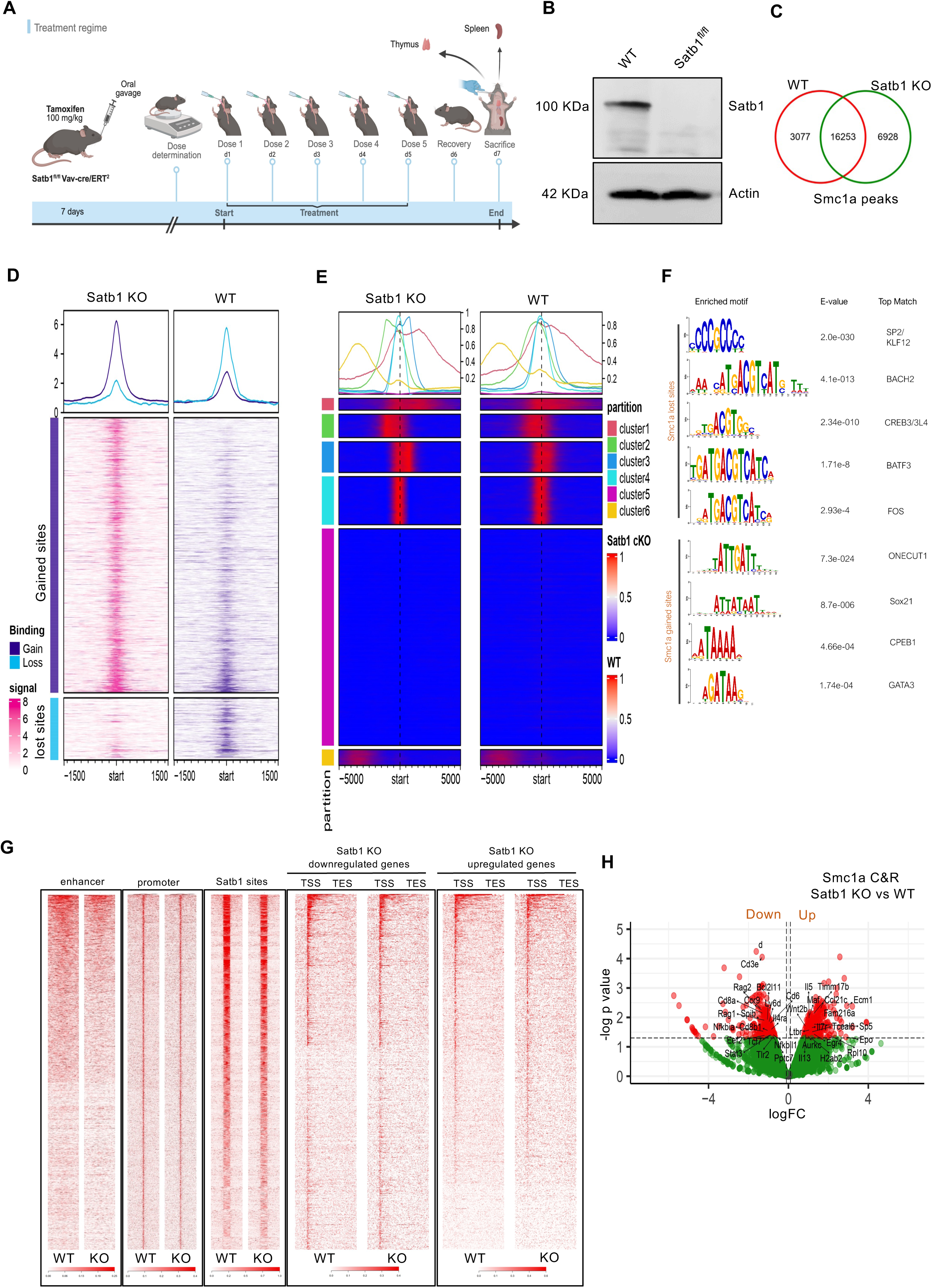
Satb1 deletion modulates chromatin binding of Smc1a in DP thymocytes. **A.** Schematic of ERT2 mediated induction to knockout Satb1 in mice. Tamoxifen induces ERT2, promoting the expression of Cre protein in Vav positive cells to delete the floxed exon-2 region in Satb1 gene. **B.** Knockout efficiency of Satb1 in DP thymocytes upon tamoxifen treatment regime depicted in **A** via western blotting. Representative of n=3 independent experiments. **C**. Smc1a Cut & Run was performed in WT and Satb1 KO DP thymocytes. Venn diagram depicting overlap of Smc1a called peaks in WT and Satb1 KO datasets. Peaks common to 2 biological replicates for each condition were used to find common and unique peaks between the sets. **D**. Differential binding analysis (using Diffbind) identified significantly enriched gained and lost peaks by Smc1a in DP thymocytes upon Satb1 depletion, shown as heatmap (2 clusters). Average profiles are shown on top of the heatmaps. **E**. k-means clustering of Smc1a peaks from WT and Satb1 KO samples. The 5 clusters are shown as heatmaps, and the overlayed average profile of each cluster is shown on top. **F**. Seq-logos of top motifs obtained from differential motif enrichment analysis in WT and Satb1 KO conditions, indicating gained and lost sequence preference. **G**. Occupancy of Smc1a on enhancers, promoters and Satb1 binding sites is shown in WT and Satb1 KO conditions. Smc1a occupancy was also plotted at TSS-TES (gene body) for downregulated and upregulated genes in Satb1 KO DP thymocytes compared to WT. **H**. Volcano plot showing the differentially bound genes by Smc1a upon Satb1 KO compared to WT DP cells. logFC of 1.5 and padj <0.05 were used as cutoff criteria.

**Figure 4.**
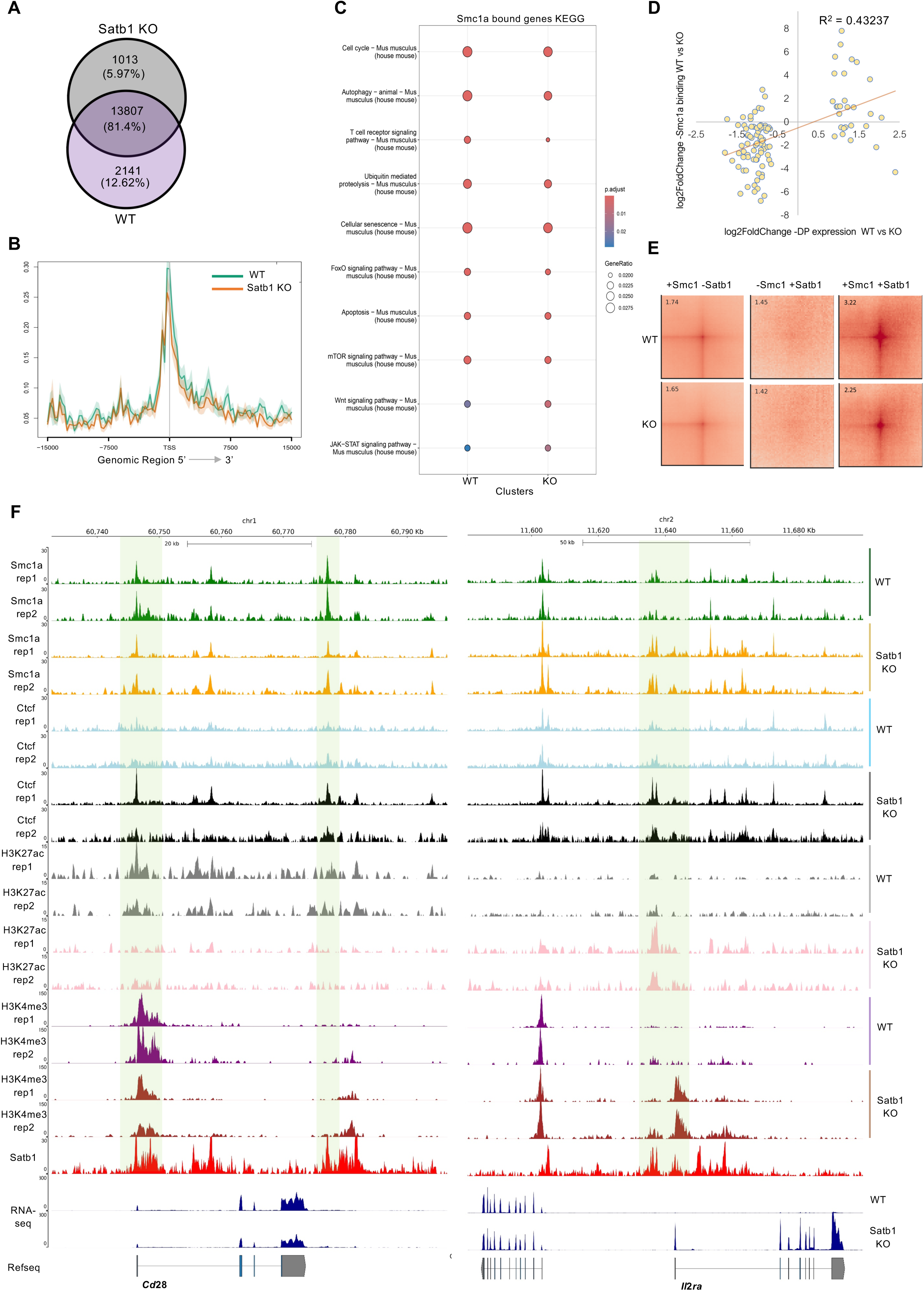
Reduction in Smc1a occupancy at T cell activation genes upon deletion of SATB1 in DP thymocytes. **A**. Venn diagram showing the ratio of common and distinct genes bound by Smc1a upon Satb1 KO in DP thymocytes. **B**. Average profile of Smc1a at TSS of T cell development associated genes in WT and Satb1 KO is shown. The gene list was obtained from publicly available data [60]. **C**. Smc1a bound genes were subjected to KEGG pathway analysis in WT and Satb1 KO conditions. The significantly enriched pathway (using BH statistic, pval <0.01) clusters in each condition were compared and shown as dotplot. **D**. Pearson correlation of genes (logFC) both differentially expressed and differentially bound by Smc1a, in WT vs Satb1 KO conditions, is shown as a scatter plot. **E**. Genomic interactions at Smc1a, Satb1 and Smc1a+Satb1 common sites are shown as pileup analysis in WT and Satb1 KO HiC samples. The score of averaged interactions (central dot) is shown at the top left corner. **F**. Genome browser tracks at *Cd28* (left) and *Il2ra* (right) loci. Individual tracks are shown as biological replicates (n=2) for Cut & Run samples. All datasets were normalized by reads per genomic content (RPGC). Corresponding gene expression in WT and Satb1 KO is shown at the bottom.

**Figure 5.**
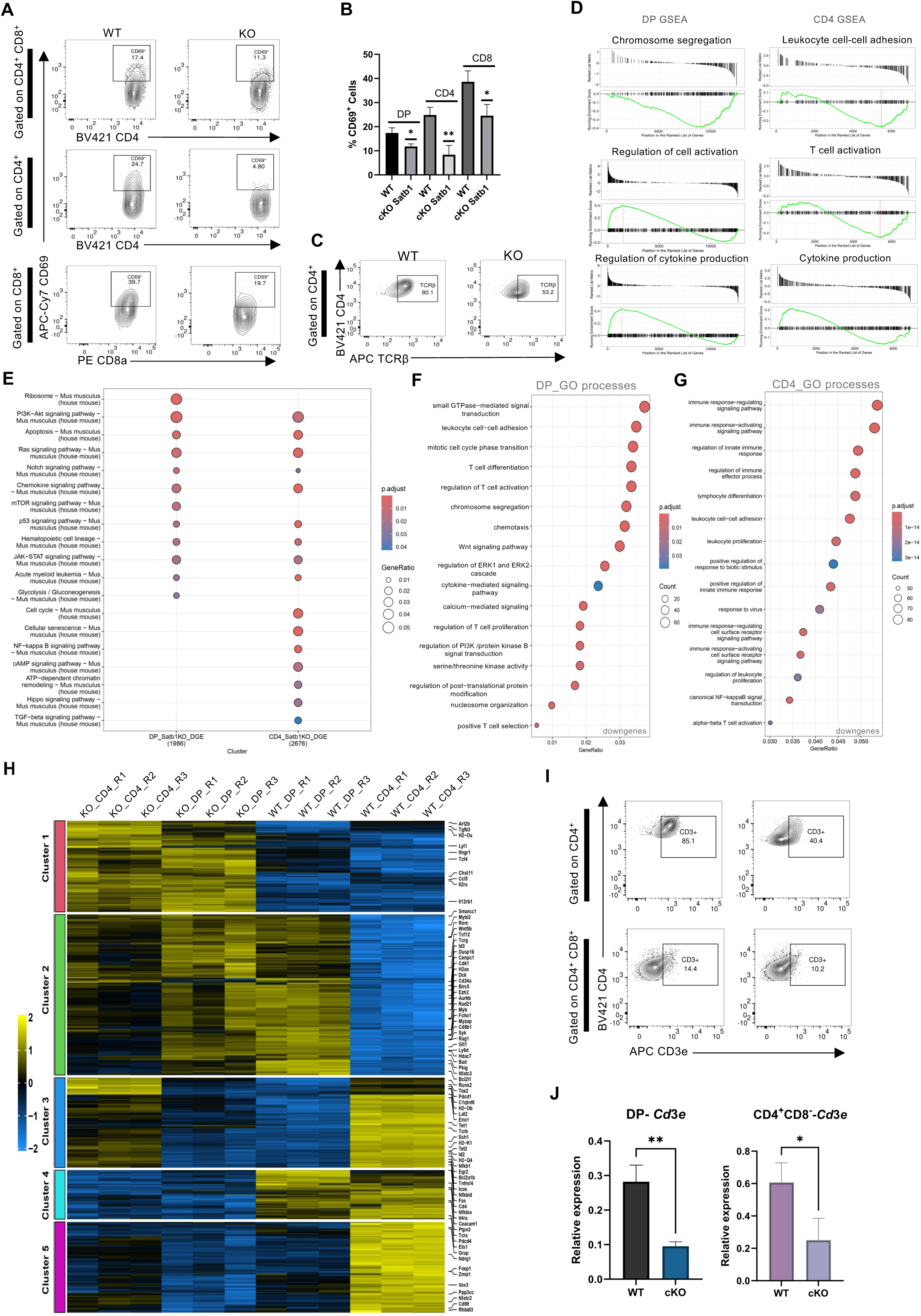
Satb1 deletion dysregulates T cell activation and cytokine signaling. **A.** Flow cytometry analysis of early T cell activation marker CD69 was performed in thymic T cells in WT and Satb1 cKO conditions. Cells were gated for DP (CD4^+^CD8^+^), CD4 (CD4^+^CD8^-^) and CD8 (CD4^-^CD8^+^). The surface expression of CD69 is shown in these populations. **B.** Quantitation of CD69^+^ cells in DP, CD4 and CD8 populations in WT and KO cells. **C.** CD4 cells were gated and TCRβ surface expression was monitored in WT and Satb1 KO thymocytes. FlowJo v10 was used for flow cytometry analysis (n=3) and population counts for CD69^+^ cells. Representative images of 3 independent replicates are shown. Graphpad v10 was used for statistical analysis using ANOVA, n=3. * p<0.05 **, p<0.01 ***, p<0.001.**D.** Gene set enrichment analysis (GSEA) of differentially expressed genes (DEGs) upon Satb1 KO in DP (left) and CD4 (right). **E**. Biological pathways, using KEGG database, for DP and CD4 DEGs upon Satb1 KO, were clustered and compared. Top enriched pathways for both datasets are shown as dotplot. **F** and **G**. Dotplots representing most significantly enriched Gene Ontology (GO) processes for downregulated genes in both DP and CD4, respectively, upon Satb1 KO. **H**. The effect of Satb1 KO on gene expression was compared for DP and CD4 thymocytes is shown as heatmap. Different clusters (k-means) show Satb1 dependent expression patterns in both cell types. **I**. CD3 surface levels in DP and CD4 SP thymocytes is shown. **J.** qRT-PCR analysis of Cd3e expression in sorted DP and CD4 thymocytes using WT controls and Satb1 KO mice. Graphpad v10 was used for statistical analysis using unpaired Student’s t-test, n=3. * p<0.05 **, p<0.01 ***, p<0.001.

**Figure 6.**
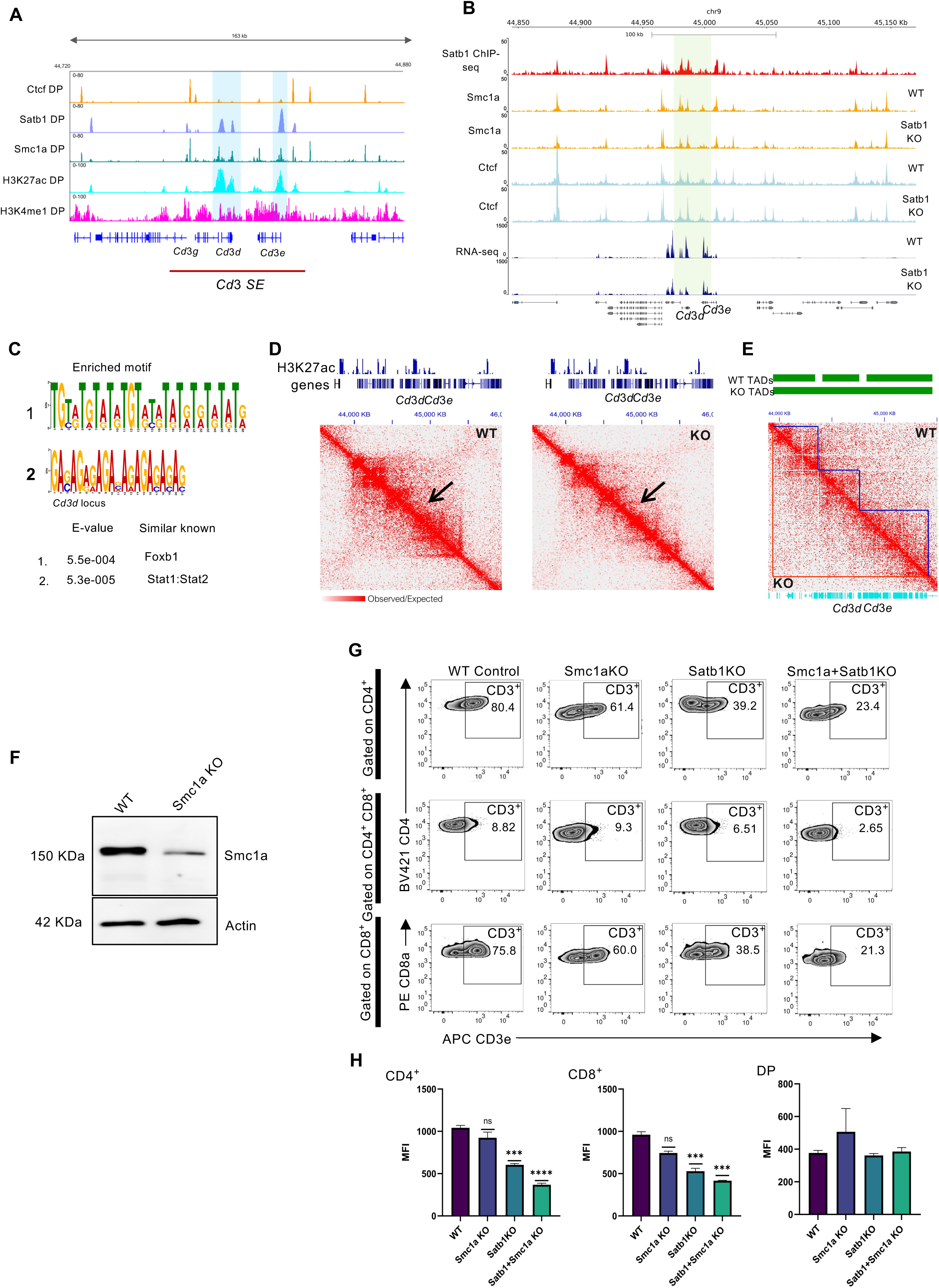
Satb1 and Smc1a co-regulate Cd3 expression in positively selected thymocytes. **A.** ChIP-seq occupancies of Ctcf, Satb1, Smc1a, as well as histone modifications H3K27ac and H3K4me1, indicating the super enhancers (SE) at the *Cd3* locus are shown as IGV tracks. **B**. Genome browser tracks at the *Cd3* locus shows binding of Smc1a and Ctcf in WT and Satb1 KO conditions. Highlighted region shows significant change in Smc1a binding in Satb1 KO compared to WT DPs. **C**. Motif enrichment analysis of 6 kb region around the *Cd3d* locus using MEME suite [51]. Significantly enriched motifs are shown. **D**. HiC data from WT and Satb1 KO in DP thymocytes is shown at 10 kb resolution. Normalization by coverage was used. Arrowheads indicate regions of reduced O/E ratios in Satb1 KO compared to WT. **E**. TADs called from WT and Satb1 KO datasets are shown (green) followed by the contact matrix in WT and Satb1 KO datasets at the *Cd3* locus. TADs are overlayed onto the matrix for visualization using Juicebox. **F**. Western blot shows efficiency of Smc1a KO using the lentiviral transduction in DP thymocytes. **G**. Flow cytometry analysis depicting CD3^+^ populations in different conditions is depicted. Representative graphs from 3 independent experiments are shown. **H**. Mean fluorescence intensity (MFI) of CD3 in different conditions as indicated. FlowJo v10 was used for flow cytometry analysis (n=3) and MFI calculations. Graphpad v10 was used for statistical analysis using ANOVA, n=3. * p<0.05 **, p<0.01 ***, p<0.001.

**Figure 7.**
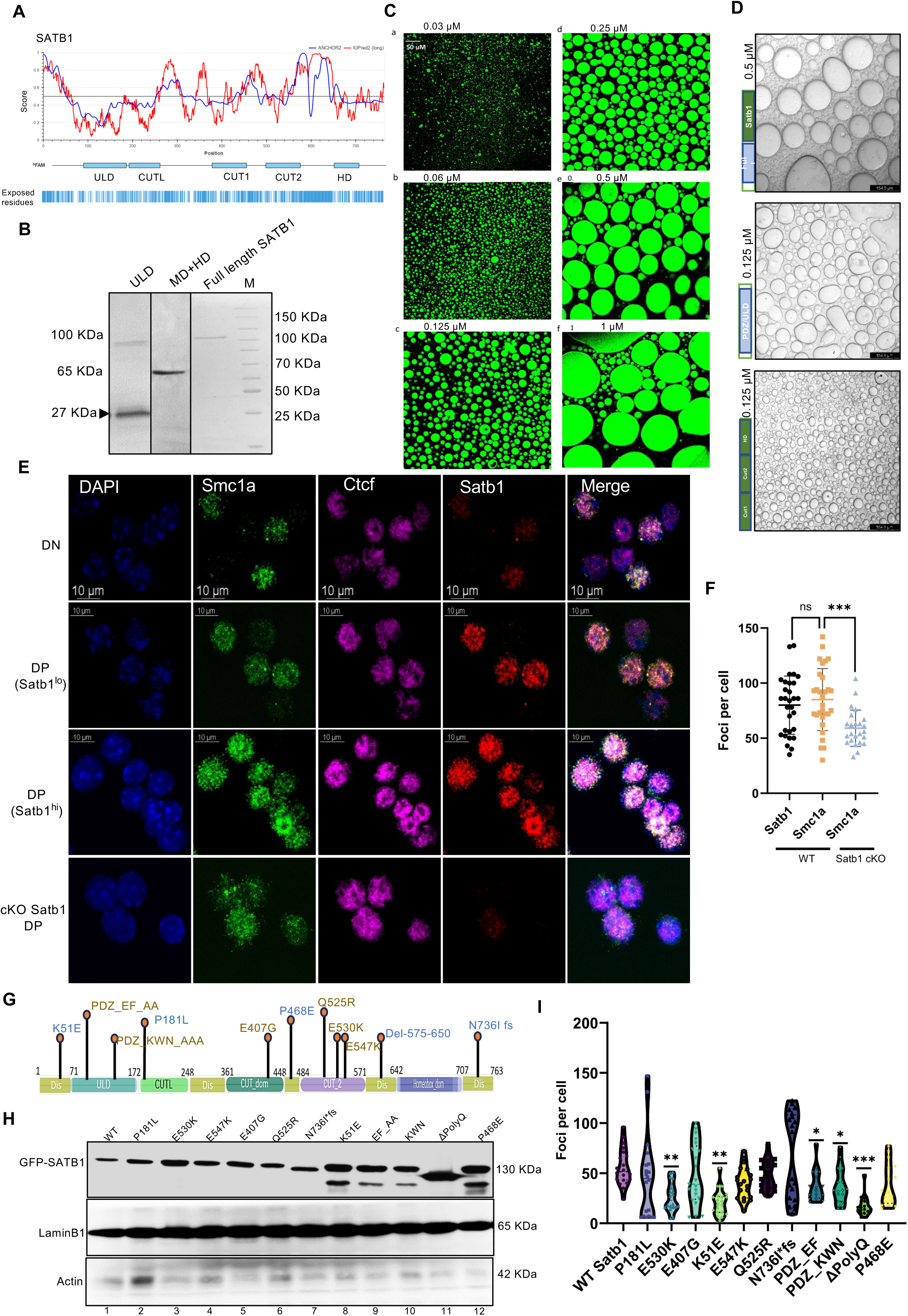
Satb1 exhibits condensate formation in vitro and in vivo. **A.** Satb1 amino acid sequence was acquired from Uniprot database (Q01826) and fed to publicly available software IUPred [94]. Scores above the baseline indicates increased disorder. **B**. Full-length SATB1 protein as well as PDZ/ULD and Cut/MAR-binding domain (MD) + homeodomain (HD) proteins were bacterially expressed and purified using Ni-NTA affinity purification. Panel depicts denaturing polyacrylamide gels (SDS-PAGE) stained with Coomassie blue. Lane ‘M’ denotes the molecular weight marker. **C**. In vitro droplet formation assay was performed with the purified Satb1 using increasing concentrations of the protein. Fluorescently labelled linearized DNA was used in the assay. The droplets containing Satb1 and bound DNA (green) were visualized using confocal microscopy. **D**. Droplet formation assay was carried out but without the presence of DNA and visualized with DIC light microscope. Purified full-length Satb1 (FL Satb1, Top), ULD (middle) and Cut1+Cut2+HD (bottom) were used for the droplet assay. **E**. Immunostaining of Satb1, Smc1a and Ctcf is shown in WT DP and DN stages is shown. Last row depicts immunostaining of these factors in Satb1 KO DPs. **F.** Quantitation of distinct foci formed by Satb1 and Smc1a in WT DP thymocytes as well as Smc1a in Satb1 KO cells. Representative images of N=3 independent experiments. Foci were counted using ImageJ, n=30. Graphpad v10 was used for statistical analysis, two-sided ANOVA was performed *** p<0.01, ns non-significant. **G.** Schematic of SATB1 domain map with its constitutive domains. The disordered regions identified are labelled ‘Dis’. SATB1 mutations identified in both structured and unstructured regions are shown. **H.** Immunoblotting shows SATB1 protein expression for all the mutant clones generated via site-directed mutagenesis. **I.** Quantitation of the number of foci formed by SATB1 mutants is shown. n=3, N=30. Graphpad v10 was used for statistical analysis, two-sided ANOVA was performed * p<0.05, ** p<0.01, *** p<0.001

### Satb1 deletion leads to repositioning of Smc1a on chromatin

We intuitively next investigated the role of Satb1 in cohesin-mediated gene regulation. We employed a tamoxifen-inducible Vav-Cre knockout system to delete Satb1 specifically from hematopoietic and T cell lineages (Figure 3A), generating Satb1^fl/fl^–Vav-iCre mice. Tamoxifen treatment resulted in >95% depletion of Satb1 in thymocytes (Figure 3B). Consistent with previous studies [23, 39] we observed a reduction in DN, CD4SP, and CD8SP populations, accompanied by an accumulation of DP thymocytes (Figure S5A). To assess the effect of Satb1 loss on cohesin binding, we performed Cut & Run analysis for Smc1a occupancy in wild-type and Satb1-deficient DP thymocytes (Figure S5B–C). Satb1 deletion led to the loss of 3,077 Smc1a peaks and the gain of 6,928 new peaks (Figure 3C–D). Clustering analysis revealed distinct changes in Smc1a binding patterns between WT and KO cells (Figure 3E), with cluster-specific shifts in average Smc1a signal upstream or downstream of peak centers in the absence of Satb1. Differential binding analysis further showed that Smc1a occupancy was significantly altered in Satb1 KO thymocytes (Figure S5D). Motif enrichment analysis of the differential peaks revealed that Smc1a binding was reduced at transcription factor motifs such as KLF12, FOS, and BATF3, while increased binding was observed at AT-rich motifs including CPEB1, ONECUT1, and GATA3 (Figure 3F). Notably, Smc1a binding was specifically reduced at enhancers and promoters, especially at Satb1-bound regulatory elements (Figure 3G and S5E). TSS-specific analysis revealed decreased Smc1a occupancy at key T cell genes such as *Cd8b1*, *Rag2*, and *Cd3d* (Figure 3H and S5F). Gene ontology analysis of WT- and KO-specific Smc1a peaks further supported the preferential loss of binding at genes involved in lymphocyte activation and immune function (Figure S5G–H). As observed, Satb1 deficiency disrupts the DP-to-SP (CD4/CD8) transition [27]; Figure S5A), we next asked whether gene expression changes in Satb1^-/-^ DPs are dependent on cohesin binding. Although ∼80% of Smc1a-bound genes in WT cells retained Smc1a occupancy in the KO condition (Figure 4A), genes involved in T cell development ([60]; Figure 4B) and T cell activation pathways showed reduced Smc1a enrichment upon Satb1 loss (Figure 4C). Furthermore, there was a positive correlation between Smc1a binding and Satb1-dependent gene expression (Figure 4D), suggesting a cooperative regulatory role between Satb1 and Smc1a during T cell development. K-means clustering of Smc1a binding sites on genes up- or downregulated in Satb1 KO DPs (Figure S6A–B) revealed specific clusters with strong correlation to Satb1-dependent transcription. We also cross-referenced dysregulated genes with Smc1a peaks gained or lost upon Satb1 depletion to pinpoint potential cohesin targets that depend on Satb1 for regulation (Figure S6C–D).

To assess the impact of Satb1 loss on higher-order chromatin architecture, we analyzed available Hi-C [39] and ChIP-seq [38] datasets from DP cells. Genomic regions co-occupied by Satb1 and Smc1a showed a marked reduction in long-range chromatin interactions in Satb1-deficient DPs (Figure 4E). While local interactions around Satb1-bound sites decreased significantly in the KO condition (Figure S7A), regions co-occupied by Ctcf and Satb1 displayed only modest changes (Figure S7B). Finally, we observed that Satb1 loss leads to redistribution of Smc1a binding at regulatory loci of T cell activation genes, accompanied by corresponding changes in gene expression and active histone modifications (H3K27ac and H3K4me3) at TCR signaling-associated genes (Figure 4F and S7C–F). Collectively, these findings suggest that in the absence of Satb1, Smc1a binding is reprogrammed—specifically diminishing at key regulatory regions—resulting in reduced chromatin interactions and impaired expression of genes critical for T cell activation.

### Satb1 depletion causes dysregulation in T cell activation and cytokine signaling in positively selected CD4 thymocytes

Given that Satb1 knockout (KO) impairs the transition from DP to SP thymocytes, we next investigated whether TCR signaling is disrupted in these cells. Effective TCR signaling is essential for DP cell survival and progression to the SP stage [61, 62]. Previous studies have implicated Satb1 in promoting CD4 SP differentiation [23, 27], in part through regulation of key genes such as *Cd4*, *Cd8*, *Runx3*, *ThPOK*, and *Foxp3* [27]. To assess the impact of Satb1 loss on positively selecting thymocytes, we measured the surface expression of CD69, an early TCR-induced marker [63] (Figure 5A). CD69 expression was significantly reduced in Satb1 KO DPs (11.3% CD69⁺) compared to WT DPs (17.4% CD69⁺) (Figure 5A). This reduction was even more pronounced in SP thymocytes: CD69⁺ CD4 cells dropped from 24.7% (WT) to 4.8% (KO), and in CD69⁺ CD8 cells from 39.7% to 19.7% (Figure 5B). A concurrent reduction in TCRβ expression further supported a defect in positive selection (Figure 5C). We then examined the transcriptional consequences of Satb1 deletion in CD4 SP versus DP thymocytes (Fig. S8A). Gene Set Enrichment Analysis (GSEA) of differentially expressed genes (KO vs WT) revealed upregulation of pathways related to cell activation and cytokine production in DPs (Figure 5D, Fig. S8B), and a marked downregulation of T cell activation and cell-cell adhesion pathways in CD4^+^ T cells (Figure 5D). While some dysregulated pathways, such as chemokine signaling and PI3K-Akt signaling, were shared between DPs and CD4s, others like cell cycle regulation and TGFβ signaling were uniquely affected in CD4 cells (Figure 5E), indicating stage-specific roles for Satb1. Gene ontology (GO) analysis identified commonly overrepresented biological processes among significantly downregulated (Figure 5F-G) and upregulated (Figure S8C-D) genes (logFC > 2) in both DPs and CD4s upon Satb1 loss. Clustering of the most dysregulated transcripts across these populations revealed overlapping and distinct gene expression changes (Figure 5H and Figure S8E-F). Notably, several TCR signaling-related genes, including *Icos*, *Cd4*, *Tcrb*, *Cd3d*, and *Nfkb*, were consistently downregulated in both populations. To further assess TCR activation, we analyzed CD3 surface expression. While CD3⁺ DP cells decreased modestly in Satb1 KO (10.2%) compared to WT (14.4%), a dramatic reduction was seen in CD3⁺ CD4 SP cells—from 85.1% in WT to 40.4% in KO (Figure 5I), consistent with the transcript-level data (Figure 5J). Overall, these findings demonstrate that Satb1 orchestrates gene expression in a stage-specific manner during T cell development, while maintaining a consistent role in regulating T cell activation and cytokine signaling across thymocyte subsets.

### Satb1 coordinates with Smc1a to ensure functional TCR signaling by regulating CD3 expression

The CD3 complex, which is non-covalently associated with the cytoplasmic tails of the TCR heterodimer, plays a critical role in transmitting TCR signals to downstream components of the signalosome cascade [62, 64, 65]. Given the observed downregulation of CD3 at both transcript and protein levels following Satb1 deletion, we investigated whether Satb1-mediated chromatin regulation contributes to CD3 expression. ChIP-seq analysis revealed that, in wild-type (WT) DP thymocytes, Satb1 binds to a super-enhancer cluster at the *Cd3* locus—marked by H3K27ac and H3K4me1—together with Smc1a [66] and Ctcf (Figure 6A), indicating potential co-regulation of the *Cd3* locus. Supporting this, Satb1 KO DP thymocytes exhibited reduced Smc1a binding at the *Cd3d* promoter and *Cd3e* enhancer (Figure 6B), while Ctcf occupancy remained largely unchanged. Motif analysis of a 5kb region surrounding the diminished Smc1a peaks in Satb1-deficient cells revealed enrichment for Foxb1 and Stat binding sites (Figure 6C), suggesting partial loss of AT-rich regions normally occupied by Smc1a. Additionally, both Ctcf and Satb1 consensus motifs were present within the *Cd3d* promoter (Figure S9A). To assess the higher-order chromatin changes, we analyzed Hi-C data from WT and Satb1 KO DPs [39] and observed a moderate reduction in intra-TAD contact frequency at the *Cd3* locus in the KO (Figure 6D). Moreover, Satb1 KO cells exhibited a TAD fusion event approximately 200 kb downstream of *Cd3*, indicating that Satb1 influences local chromatin architecture essential for proper CD3 expression (Figure 6E). Given the loss of Smc1a binding in the absence of Satb1, we further explored whether Smc1a and Satb1 cooperatively regulate CD3. Using CRISPR/Cas9 lentiviral delivery, we depleted Smc1a in thymocytes (Figure 6F) and assessed CD3 expression across thymocyte subsets by flow cytometry (Figure 6G). Smc1a knockout alone caused only a modest reduction in CD3^+^ CD4SP (19%) and CD8SP (15%) cells relative to control (Figure 6H). In contrast, Satb1 knockout, as previously shown (Figure 5I), resulted in a substantial decline in CD3^+^ SP populations. Notably, co-deletion of Smc1a and Satb1 led to a more pronounced decrease in CD3^+^ cells (Figures 7H and S9B), indicating a synergistic regulatory role. To validate their cooperative function, we performed a dual-luciferase reporter assay using a known Satb1 target promoter construct (Figure S9C) [31]. Co-expression of Satb1 and Smc1a significantly enhanced luciferase activity compared to either factor alone (Figure S9D), further supporting their interaction.

Chromatin accessibility dynamically shifts during T cell development to meet stage-specific transcriptional demands [52]. At the *Cd3* locus, chromatin becomes accessible starting from the DN2b stage (Figure S10A). We examined whether Satb1 deletion alters this accessibility using ATAC-seq in WT and Satb1 KO DP cells [39]. The results showed decreased accessibility at the *Cd3* locus in KO cells (Figure S10B), while histone modifications such as H3K4me3 and H3K27me3 remained unchanged (Figure S10C). Furthermore, ATAC-seq across four developmental stages (DN, DP, CD4, CD8) revealed a mild reduction in *Cd3e* promoter accessibility at DN and DP stages (Figure S10D), but a substantial (∼6-fold) loss of accessibility in Satb1 KO CD4 thymocytes compared to WT (Figure S10D). Together, these findings demonstrate that Satb1 regulates CD3 expression during DP-to-SP transition by coordinating chromatin interactions through its cooperation with cohesin and by modulating chromatin accessibility in a stage-specific manner.

### Satb1 undergoes liquid–liquid phase separation to orchestrate transcriptional regulation

Since Satb1-mediated genomic interactions—particularly in coordination with cohesin (Smc1a)—are essential for maintaining the transcriptional program during the DP to SP transition in T cells, we next explored whether specific biophysical properties of Satb1 contribute to its function. We observed that Satb1 forms distinct speckle-like nuclear foci in DP thymocytes (Figure 2C), indicative of potential condensate formation [67] This promted us to investigate whether Satb1 exhibits characteristics of liquid–liquid phase separation (LLPS). *In silico* analysis of the Satb1 primary amino acid sequence revealed several intrinsically disordered regions (IDRs) interspersed between structured domains (Figures 7A and S11A), a pattern also confirmed by *de novo* structural predictions (Figure S11B). Unlike other transcription factors known to undergo LLPS, such as TCF1 and BRD4, which often contain continuous IDR stretches, Satb1 presents a more segmented IDR profile (Figure S11C). Structural modeling of Satb1 dimers and trimers further suggested that its IDRs may interact intermolecularly, in addition to dimerization via the ULD (Figure S11D). To test Satb1’s LLPS propensity in vitro, we performed droplet formation assays using purified full-length Satb1 protein and its functional domains (Figure 7B, schematically depicted in Fig. S4A). Satb1 readily formed liquid-like droplets in the presence of DNA, with droplet size scaling with protein concentration (Figure 7C), and was also capable of phase separating in the absence of DNA (Figure 7D). To determine the domain contributions to condensate formation, we expressed and purified two functional regions—ULD/PDZ and the DNA-binding Cut1+Cut2+HD (MD+HD)—and found that both domains independently formed droplets (Figure 7D), consistent with their inclusion of IDRs. These condensates exhibited hallmark liquid-like properties, including dynamic molecular exchange as shown by fluorescence recovery after photobleaching (FRAP) (Figure S12A-B). Satb1 droplets were also capable of sequestering DNA in a concentration-dependent manner (Figure S12C). However, as PEG was required to observe droplets and PEG is not always an inert crowding agent, we note there are limitations in making strong conclusions about the phase-separation properties of Satb1 [68–70].

Furthermore, treatment with 1,6-hexanediol (1,6-HD), known to disrupt weak hydrophobic interactions underlying LLPS [71], resulted in droplet dissolution (Figure S12D), reinforcing the liquid-like nature of Satb1 condensates. To assess whether these phase separation properties relate to Satb1’s molecular interactions in T cell development, we examined its nuclear distribution in DP thymocytes. Satb1 appeared in discrete foci (Figure 7E), while Ctcf showed diffuse nuclear staining. Interestingly, Smc1a formed distinct nuclear speckles that co-localized with Satb1 foci, consistent with its predicted disordered regions (Figure S13A-D). Notably, Satb1 KO DPs showed reduced Smc1a foci (Figure 7F), suggesting that Satb1 contributes to cohesin enrichment in nuclear condensates. Disruption of LLPS by 1,6-HD led to significant loss of both Satb1 and Smc1a foci, which partially recovered upon alcohol washout (Figure S13E-F). Given that 1,6-HD has also been implicated in non-specific effects in the nucleus, we interpret these cellular results with 1,6-HD conservatively [72]. However, together, the findings suggest that Satb1 has a propensity for LLPS and may function by forming nuclear condensates that facilitate recruitment of Smc1a during T cell development.

Recent studies have linked specific mutations in the SATB1 coding sequence to a range of neurodevelopmental disorders, including intellectual disability, dental anomalies, epilepsy, behavioral disturbances, and facial dysmorphisms [73]. In addition, a premature truncating mutation in SATB1 has been identified in lung cancer [74], and another report described a pathogenic SATB1 variant in a case of Trisomy X syndrome [75]. Given the presence of intrinsically disordered regions (IDRs) throughout the SATB1 protein, we hypothesized that these single nucleotide polymorphisms (SNPs) might disrupt its ability to form condensates. To test this, we selected reported mutations located in both structured domains and IDRs of SATB1 (Figure 7G) and introduced them into a SATB1-GFP expression construct using site-directed mutagenesis. Proper nuclear expression of each mutant was validated by overexpression in HEK293T cells (Figure 7H). We then assessed their impact on condensate formation via live-cell imaging of transfected HEK293T cells (Figure S14A). Compared to wild-type (WT) SATB1, the K51E mutation in the ULD-associated IDR and the ΔPolyQ mutation in the C-terminal IDR both showed reduced capacity for phase separation (Figures 7I and S14A). Similarly, mutations in structured regions—E530K in the CUT domain, and EF→AA and KWN→AAA in the ULD domain—also exhibited a significant loss of nuclear foci (Figure 7I). To evaluate the functional consequences of impaired condensate formation, we performed a dual-luciferase reporter assay using the *Il2* promoter to measure the transcriptional activity of these SATB1 point mutants. Notably, SATB1 mutants with compromised phase separation potential exhibited decreased reporter expression compared to WT (Figure S14B). Together, these findings indicate that specific amino acid substitutions in SATB1 can impair its phase separation behavior, leading to dysregulation of its transcriptional targets.

## Discussion

In this study, we uncover a nuanced regulatory function of Satb1 during T cell development, operating through its association with Ctcf and cohesin complex. While Ctcf and cohesin are well-characterized for their roles in genome looping, TAD formation, and global chromatin architecture across various cell types [1], our data suggest that Satb1 co-occupies Ctcf- and cohesin-bound genomic sites in DP thymocytes, frequently overlapping with enhancer regions. These co-bound regions show increased chromatin interactions as T cell development progresses. Moreover, Satb1 localizes to TAD boundaries alongside Ctcf and is enriched at regulatory loci of boundary-associated genes, suggesting that it may influence inter- and intra-TAD interactions. Distinct from Ctcf, Satb1 expression is specifically upregulated in T cells, particularly from the DP stage onward, hinting at a dynamic, TCR signal-responsive role in shaping chromatin architecture during thymic development. Cooperative actions among lineage-defining transcription factors are common in T cells—for instance, TCF1 recruits HEB1 to modulate the DP-specific gene program by influencing traditional and super-enhancer activity [76].

Satb1 interacts with a diverse array of proteins, including histone modifiers such as HDACs [31] and p300 [19]), chromatin remodelers like SWI/SNF and NuRD [33, 77], as well as transcription factors such as β-catenin. Our mass spectrometry analysis revealed DP-stage-specific Satb1 interactors involved in chromatin remodeling, translation, and chromatin modification. Among these, we validated Smc1a, a core subunit of the cohesin complex, as a direct interactor. Cohesin is well known for its roles in chromatin loop extrusion, cell cycle progression, and chromatin condensation [78].

Previous studies examined Satb1 function in T cells via conditional knockouts using Lck- or CD4-Cre drivers [23, 28] or pan-hematopoietic Vav-Cre [39]. To study its function in a temporally controlled manner, we employed an inducible Vav-CreERT2 Satb1 knockout model, which recapitulated previously observed thymic defects. In the absence of Satb1, Smc1a chromatin occupancy was disrupted at Satb1-bound and immune-regulatory loci, such as Cd8b1 and Cd3d, demonstrating Satb1-dependent recruitment of cohesin at specific genomic sites.

Satb1 deletion impairs differentiation from DP to single-positive (SP) CD4 and CD8 T cells and affects regulatory T cell (Treg) development [23, 27]. Our findings indicate that Satb1 regulates downstream TCR-responsive genes—including CD69 and TCRβ—during DP-to-SP transition, underscoring its role in positive selection. Interestingly, although changes are moderate in DPs, CD4 and CD8 SP thymocytes exhibit a more profound dysregulation, suggesting a sustained role for Satb1 in maintaining TCR-mediated activation post-differentiation.

We further identified a critical function of Satb1 in regulating CD3 surface expression in SP thymocytes, which correlates with altered chromatin organization at the *Cd3* locus in Satb1-deficient cells. This may explain the reduced responsiveness of Satb1-deficient DP thymocytes to anti-CD3/CD28 stimulation [23]. Satb1 is known to bind nucleosomes strongly [79], indicating its association with closed chromatin. At the *Cd3* locus, Satb1 contributes to maintaining an open chromatin state across multiple developmental stages, with a more pronounced effect in CD4 SPs than in DPs. Additionally, we show that Satb1 orchestrates transcriptional programs related to T cell activation and cytokine signaling in both DPs and CD4 SPs.

Emerging evidence implicates biomolecular condensates in organizing the genome [80] and regulating transcription [81]. More recently, a long form of Satb1 (Satb1L) was shown to phase-separate which was dependent on post-translational modifications [82]. Our data suggest that Satb1 forms nuclear condensates, facilitated by its intrinsically disordered regions (IDRs), which are distributed across its ULD and MD+HD domains. These flexible regions may enable Satb1 multimerization via multiple interfaces. We found that in vitro phase separation by Satb1 is enhanced in the presence of DNA—though not strictly required for demixing, DNA promotes more functional condensate formation. In the absence of DNA, higher Satb1 concentrations are needed, and the resulting droplets display irregular morphology. Thus, while dispensable for condensate formation, DNA binding appears critical for modulating the biophysical and functional properties of Satb1 condensates. The cohesin complex and Ctcf are key regulators of chromatin architecture, maintaining genome organization through long-range chromatin interactions [10]. Recently, biomolecular condensates have emerged as a crucial link between chromatin structure and gene function, offering new insight into genome organization [67, 81, 83]. Satb1, a chromatin organizer, serves as a scaffold for recruiting chromatin modifiers and nucleosome remodelers, thereby influencing the transcriptional activity of its target genes [33]. Satb1 anchors DNA to the BURs in the nuclear matrix [84] mediating chromatin looping at loci such as Th2 cytokine genes [32] and the MHC-I locus [38]. Our findings suggest that Satb1 co-occupies chromatin with Smc1a and Ctcf, and that these interactions may be, at least in part, mediated by phase separation. Notably, Satb1 depletion led to reduced nuclear foci formation by Smc1a, indicating that heterotypic interactions between Satb1 and Smc1a may facilitate condensate formation at shared genomic sites.

Liquid-liquid phase separation (LLPS) has been increasingly linked to disease pathogenesis. Several studies have highlighted how specific amino acid residues drive phase separation, and how their mutation can disturb cellular homeostasis, contributing to disease [85, 86]. For instance, many mutations associated with amyotrophic lateral sclerosis (ALS) impair LLPS, leading to neurodegeneration through loss of protein dynamics and formation of static aggregates [87]. However, the molecular mechanisms by which individual mutations influence LLPS remain incompletely understood. Some mutations reduce the saturation concentration for phase separation, promoting irreversible aggregation, while others may prevent condensate formation entirely. Our results demonstrate that disease-associated mutations in Satb1 can impair its phase separation capacity, with mutations in both IDRs and structured domains diminishing its ability to undergo LLPS. This reduction in phase separation correlates with a loss of Satb1’s gene regulatory function, underscoring the importance of homotypic interactions mediated by Satb1’s ULD domain and π–π interactions driven by its IDRs in condensate formation.

In conclusion, our findings demonstrate that Satb1 acts as a chromatin architectural protein during T cell development, functioning in concert with Ctcf and Smc1a. Satb1 physically interacts with Smc1a and may guide its chromatin association via sequence-specific recruitment, thereby modulating cohesin-dependent chromatin looping. Moreover, Satb1 and Smc1a co-regulate gene expression at a subset of common genomic loci. The ability of Satb1 to undergo phase separation further reinforces its architectural role and influences Smc1a chromatin occupancy and downstream transcriptional programs.

## Materials and Methods

### Mice

Six- to ten-week-old WT Satb1 fl/fl (Gift from Dr Terumi Kohwi-Shigematsu) and VaviCreERT2 Satb1 fl/fl mice were used to prepare the single cell suspension of thymocytes for the flow cytometry analysis as well as for sorting the subpopulations of thymocytes which are in different stages of T-cell development. All the strains were bred and maintained in sterile environment and the dissections were carried out in accordance with the guidelines of NFGFHD, IISER Pune and CITRES, Shiv Nadar University.

### Flow cytometry

The suspension of thymocytes and or splenocytes were prepared using thymii from 6-week-old mice from various genotypes indicated and were used for surface staining. Before staining, Fc -blocking was done with anti-CD16.32 antibody (BD Biosciences). Then, the thymocytes were subjected to surface staining: BV 421 anti-mouse CD4 (Clone GK1.5, Biolegend); APC anti-mouse CD3 (Clone GK1.5, eBioscience); PE-Cy7 anti-mouse CD69 (Clone GK1.3, BD Biosciences); PE anti-mouse CD8a (Clone 53-6.7, BD Biosciences); FITC TCRβ (Clone M1/69, eBioscience). For Splenic cells, APC anti-mouse CD62L (Clone 19.3, eBioscience) and PE-Cy7 CD44 (Clone B21.1, Biolegend) were used in combination with CD4 and CD8a staining. The analysis was performed in BD Celesta or BD Aria III Fusion SORP. The thymocyte sub populations such as CD4-CD8-double negative (DN), CD4^+^CD8^+^ double positive (DP), CD4^+^ SP thymocytes (total CD4SP), and CD8^+^ SP thymocytes (total CD8SP) were FACS sorted using FACS Aria III SORP (BD biosciences). The splenic CD4 subpopulations; activated and naïve were sorted based on CD44 and CD62L markers using FACS Aria III SORP (BD Biosciences).

### Datasets and bioinformatics analysis

ChIP-seq studies of the genome-wide occupancy of H3K4me3, H3K27me3, and H3K4me1 in mouse DP thymocytes were conducted using accession number GSE20898. Datasets for Satb1, Smc1a and Ctcf ChIP seq were obtained from accession numbers GSM1617950, GSE41743 and GSE61428. For HiC in Satb1 WT and cKO conditions, we used the datasets from GSE173446. Bowtie2 was used to align the raw readings, and MACS2 was used to call peaks. The IGV genome browser was used to see the peaks that MACS created. R software ChIPseeker was used to plot genome-wide occupancy profiles and biological processes. The GSE48138 dataset was used to conduct RNA-seq analysis on DP and CD4SP thymocytes. Datasets from GSE182995 was used for ATAC-seq and HiC analysis as well as for H3K4me3 and H3K27me3 modifications; both in WT and Satb1 cKO conditions. GSE77695 data set was used for the analysis of ATAC-seq performed in HSCs, MPPs and CLPs. GSE100738 data set was used for ATAC-seq analysis performed in DN1, DN2a, DN2b, DN3, DN4, DP, CD4SP and CD8SP. IGV v2.1 was used to plot the occupancies of ChIP-seq and ATAC-seq processed data onto selected genomic regions. HISAT2 and Stringtie were used to align RNA-seq reads. Additional differential gene expression analysis was conducted using R tool Deseq2.

### Cloning

Satb1 and its specific domains PDZ/ULD and MD+HD were cloned into pTRIEX-3-neo to create C-terminal HIS tag as a fusion ORF with each domain. Full length Smc1a and its identified domains N-term, Coil-I, Hinge, Coil-II and C-term were cloned in pGEX4T1 upstream of the glutathione S transferase (GST). For CRISPR, gRNA sequences for Smc1a and Satb1 were obtained by using Benchling CRISPR tool (https://www.benchling.com/crispr). The top hits were added with Bsmb1 restriction sites and used for oligo synthesis. The complimentary oligos for each gRNA were annealed together at 95°C for 5 min and cooled slowly till 25°C. The annealed oligo was ligated into linearized (with BsmbI) LentiCRISPR v2 (a gift from Feng Zhang, Addgene plasmid # 52961).

### Cell culture, transfection and lentivirus production

Growth media supplemented with 10% fetal bovine serum (Gibco) was used to cultivate HEK 293T cells, which were then kept in a humidified incubator at 37°C and 5% CO_2_. Transfection of packaging vectors pMD2 and pPAX2, along with lentiCRISPR v2 cloned with individual gRNA was performed using Lipofectamine 3000 (Invitrogen). After 48 and 72 hrs the virus containing media was collected and filtered using 0.45 μm syringe filter and stored in -80°C for future use.

### Lentiviral knockout of Satb1 and Smc1a in DP thymocytes

To isolate the thymus, C57BL/6 mice from 6 weeks old were used. A single cell suspension was made by crushing the thymus through a 70-micron filter. The following fluorochrome-tagged antibodies were then used to surface stain thymocytes: PE anti-mouse CD8a (Clone 53-6.7, BD Biosciences) and FITC anti-mouse CD4 (Clone GK1.5, BD Biosciences). Thymocytes that were double positive (DP) and CD4^+^CD8^+^ were sorted using the FACS Aria III SORP (BD Biosciences). To the sorted DP, lentivirus particles containing empty vector, Satb1 gRNA, Smc1a gRNA, or Satb1 and Smc1a gRNA were introduced. For 36 h, thymocytes were grown in RPMI-1640 medium supplemented with 10% FBS, penicillin-streptomycin, and IL7 at 5 ng/mL, SCF at 5 ng/mL, and IL2 at 2 ng/mL. Following incubation, cells were taken out and subjected to flow cytometry analysis, western blotting or RNA isolation.

### CUT&RUN sequencing

DP thymocytes from WT and Satb1 KO mice were used for CUT&RUN using the benchtop protocol [88]. One million cells were sorted and washed in ice-cold phosphate-buffered saline (PBS). Nuclei were isolated by hypotonic lysis in 1 ml NE1 (20 mM HEPES-KOH pH 7.9; 10 mM KCl; 1 mM MgCl_2_; 0.1% Triton X-100; 20% Glycerol) for 5 min on ice followed by centrifugation as above. Nuclei were briefly washed in 1.5 ml Buffer 1 (20 mM HEPES pH 7.5; 150 mM NaCl; 2 mM EDTA; 0.5 mM Spermidine; 0.1% BSA) and were incubated with pre-washed Concanavalin A magnetic beads on RT for 5 min. Nuclei-bound beads were washed in 1.5 ml Buffer 2 (20 mM HEPES pH 7.5; 150 mM NaCl; 0.5 mM Spermidine; 0.1% BSA). Nuclei were resuspended in 500 µl Buffer 2 and 10 µl SMC1A antibody (Abcam) or CTCF (Cell Signaling Technologies) was added and incubated at 4°C for 2 hr. Nuclei were washed 3 x in 1 ml Buffer 2 to remove unbound antibody. Nuclei were resuspended in 300 µl Buffer 2 and 5 µl pAG-MNase (EpiCypher) added and incubated at 4°C for 1 hr. Nuclei were washed 3 x in 0.5 ml Buffer 2 to remove unbound pA-MN. Tubes were placed in a metal block in ice-water and quickly mixed with 100 mM CaCl_2_ to a final concentration of 2 mM. The reaction was quenched by the addition of EDTA and EGTA to a final concentration of 10 mM and 20 mM respectively. Cleaved fragments were liberated into the supernatant by incubating the nuclei at 4°C for 1 h, and nuclei were pelleted by centrifugation as above. DNA fragments were extracted from the supernatant and used for the construction of sequencing libraries.

The NEB-Next UltraII library prep kit (NEB) was utilized to prepare a library with an equivalent quantity of DNA (∼5 ng) as input. Quantitative PCR (qPCR) analysis was used to estimate the number of cycles required for the amplification of adapter-ligated libraries. Using the HiPrep PCR clean up system (MagBio Genomics, USA), final libraries were purified. Prior to pooling libraries at an equimolar ratio, the library concentration was ascertained using Qubit, and the average fragment size was measured using the dsDNA HS test on the Bioanalyzer 2100 (Agilent). The HiseqX platform was utilized to acquire sequencing reads (150 bp PE).

### RNA-seq

RNA was extracted from 1-3 million sorted cells using TRIzol (Thermo F isher Scientific) for bulk RNA-seq. The Agilent 2100 Bioanalyzer was used to quantify and qualify the total RNA. Following this, 1 μg of total RNA was utilized to prepare the library. The poly(A) mRNA magnetic isolation module was used to carry out the poly(A) mRNA isolation. First strand synthesis mix (NEB) was used to create first-strand cDNA, while second strand synthesis enzyme mix (NEB) was used to create second-strand cDNA. Three steps were taken with the isolated double-stranded cDNA: adapter ligation, 3’-dA tailing, and end repair.

The number of PCR cycles was determined by qPCR. Ligated DNA was amplified using six to ten cycles with Illumina P5/P7 primers. PCR products were purified using magnetic beads and quantified using a Qubit 4.0 Fluorometer (Thermo Fisher Scientific). The resulting libraries were pooled and sequenced on the Illumina HiSeqX platform using 2×150 bp paired-end reads.

### cDNA synthesis and Quantitative PCR analysis (qRT-PCR)

RNA isolation from DP thymocytes sorted from WT and Satb1 KO mice was performed using RNeasy mini kit (Qiagen). After DNase I (Roche) treatment, the RNA was cDNA synthesis was done using High-capacity cDNA synthesis kit (Applied Biosystems). Quantitative PCR was carried out with qPCR master mix (Takara) using the following PCR conditions: 95°C-5 min, 95°C-30 sec, 60°C-30 sec, 72°C-45 sec - for 37 cycles, 72°C-3min. The primers used for RT-qPCR are shown in Table S1.

### Assay for transposase-accessible chromatin (ATAC) -qPCR

To prepare for ATAC reaction, 50,000 cells were washed twice with PBS, lysed in 50 μL of ice-cold Lysis Buffer (50 mM HEPES, 0.1% NP40,50 mM NaCl). The isolated nuclei were then pelleted for ten minutes at 500 X g at 4°C. 100 nM Tn5 Transposase was used to execute the tagmentation process in 1X Tagmentation Buffer (10 mM Tris pH 7.4, 5 mM MgCl_2_, 10% DMF, 33% PBS, 0.1% TWEEN 20, 0.01% Digitonin) for 30 min at 37°C. DNA was purified using PCR purification columns (MN biosciences). Before qPCR, DNA was eluted into a volume of 20 μl and kept at -20°C. After amplification using one set of i5 and i7 primers (NExtera), qPCRs were performed for *Cd3e* promoter using the promoter-targetting primers (Table S2).

### Co-Immunoprecipitation

Single-cell suspension of thymocytes was prepared from thymi isolated from six-week-old mice. BCA kit (Thermo Fisher Scientitic) was used to assess the protein content after cells were lysed using NP40-based lysis buffer. Using 3 μg mouse or rabbit IgG and protein A/G Dyna beads (Thermo Fisher Scientific), 500 μg of protein was precleared for 1 h. Using anti-SATB1 antibody, the cleared lysate was subjected to antibody-binding for 4 h at 4°C. Following incubation, protein A/G magnetic beads (Thermo Fisher Scientific) were used to precipitate anti-SATB1-bound protein complexes. The beads were washed three times with lysis buffer. Bound proteins were eluted using 1% SDS in 1X PBS at 98°C for 15 min. Co-immunoprecipitated proteins were identified by subjecting the eluted protein to SDS gel electrophoresis followed by western blotting using anti-SMC1A (Abcam).

### Mass spectrometry

For mass-spectrometry samples, the protein A/G beads with immunoprecipitated proteins were washed 3 times with 1x PBS. 2M urea prepared in triethylammonium bicarbonate (100 mM) was added to the washed beads. 200 mM Tris(2-carboxyethyl)phosphine (Sigma-Aldrich) was added, and the samples were incubated at 37°C for 30 min at 500 rpm. 200 mM of iodoacetamide (Sigma-Aldrich) was added to the mixture and incubated at RT for 30 min in dark with intermittent mixing. MS-grade trypsin (Promega) was added at a concentration of 1μg/μl and the samples were incubated at 37°C overnight. Trifluoroacetic acid (Sigma-Aldrich) was added to final concentration of 0.1 percent to quench trypsin activity. Desalting of digested samples was carried out using C18 ziptips (Millipore). The C18 column was activated using 100% acetonitrile buffer (ACN) and was equilibrated using 0.1% TFA. The samples were loaded to the ziptips and flow-through was discarded. The columns were washed 5x with 0.1% TFA, followed by elution with 60% ACN prepared in 0.1% TFA. The collected flow-through was dried and stored at -20°C for sample preparation. LC-MS was performed at IISER Pune Mass Spectrometry facility, and the data was analyzed using ProteinPilot (Sciex) using mouse proteome as reference to obtain the significant protein list. The significantly enriched proteins were subjected to cytoscape analysis for pathways and network (PPI) enrichment.

### Western blotting

Protein quantification was performed using the BCA assay kit (Thermo Fisher Scientific) following cell lysis with RIPA buffer (50 mM Tris-HCl, pH 7.4, 150 mM NaCl, 1 mM EDTA, 0.5% NP-40, 0.25% sodium deoxycholate, 1 mM PMSF, and 1X protease inhibitor cocktail (Roche)). The lysates were separated on a 10% SDS-PAGE gel and transferred onto a PVDF membrane (Millipore). To block non-specific binding, the membrane was incubated in 5% skimmed milk in TBST. Immunoblotting was performed using the following primary antibodies: anti-Actin (1:1000, Millipore), anti-SATB1 (1:1000, Santa Cruz Biotechnology), and anti-Smc1a (1:1000), incubated overnight at 4°C. The next day, membranes were washed five times with 1X TBST (500 mM NaCl, 20 mM Tris pH 7.5, 1 mM EDTA, 1% Tween-20) and probed with HRP-conjugated secondary antibodies (anti-mouse IgG-HRP or anti-rabbit IgG-HRP). After additional washes with TBST, signals were developed using ECL chemiluminescent substrate (Bio-Rad) and visualized with the ImageQuant LAS 4000 system (GE Healthcare Life Sciences).

### HiC and HiChIP analysis

After adapter cutting and quality filtering, reads were acquired using fastp. Hi-C mapping was carried out using HiC-Pro. The mm10 mouse reference genome was mapped to the paired end reads. HiC-Pro was used to filter valid reads after reference mapping. At various resolutions such as 10-kb, 100-kb, and 1 Mb, raw contact matrices were created. HiC-Pro (v2.11.1) was employed to the raw contact matrices from Hi-C data. The .hic files were then generated using HiC-Pro. Using Juicebox, Hi-C contact matrices were visualized. Similar processing was used for HiChIP data. HiChIP data was plotted using Hicexplorer [89].

For pileup analyses, we utilized coolpup.py [90]. ChIP-seq peaks obtained from MACS were used to plot the interacting regions. The padding was set to 500kb for local pileup. For the aggregate analysis, interactions from 100kb to 600kb was averaged. We used different groups; Ctcf only, Satb1 only, Ctcf+Satb1, Smc only and Smc+Satb1 obtained using bedtools [91] intersect with -wa -f 0.6.

### Immunostaining

Single-cell suspensions were prepared from the thymi of six-week-old WT and Satb1 cKO mice. Thymocytes were fixed using 2% paraformaldehyde and permeabilized with 0.1% Triton X-100. Cells were then incubated for 3 h at room temperature with primary antibodies: mouse anti-SATB1 (Abcam), goat anti-CTCF (Cell Signaling Technologies), and rabbit anti-SMC1A (Abcam). Following intracellular staining, cells were washed with 1X PBS containing 0.01% Tween 20. Cells were then incubated with fluorochrome-conjugated secondary antibodies (anti-mouse GFP 488, anti-goat Alexa Fluor 568, and anti-rabbit Alexa Fluor 594) for 1 h at room temperature. DNA was subsequently stained with DAPI (Sigma). Imaging, including individual and Z-stack captures, was performed using a Leica SP8 confocal microscope.

### Proximity ligation assay (PLA) for Smc1a and Satb1

Total thymocytes and MACS-sorted CD4^+^ T cells were fixed with 4% PFA for 30 min at room temperature (RT) and permeabilised using 0.5% Triton X-100 at RT for 30 min. The PLA was performed essentially as described [92]. Blocking was done in a drop of blocking solution (Sigma-Aldrich #DUO92002) at 37 °C for 1h. Cells were incubated with rabbit anti-SATB1 (mAb 1:200; Abcam #AB109122) and mouse anti-SMC1a (mAb 1:100; CST#6892S) primary antibodies overnight at 4°C (diluted in Duolink Antibody Diluent, Sigma-Aldrich #DUO92002). Next, cells were washed twice using 1X wash buffer A, followed by incubation with PLUS (Anti-Rabbit) and MINUS (Anti-Mouse) (Sigma-Aldrich #DUO92002 & #DUO92004) probes for 1 h at 37 °C. Cells were again washed twice in wash buffer A and then incubated with the ligation mixture (ligase diluted 1:40 in ligation buffer) (Duolink In Situ Detection Reagents Orange kit, Sigma #DUO92007) for 30 min at 37 °C. Following the next round of washing in wash buffer A, cells were incubated with the amplification solution (polymerase diluted 1:40 in an amplification buffer) for 100 min at 37 °C. Finally, the cells were washed twice in 1X wash buffer B and once in 0.01X wash buffer B. Mounting was done using mounting media with DAPI (Sigma-Aldrich #DUO82040) and allowed to dry before imaging. Negative controls were performed using only the secondary antibodies. Images were acquired using a confocal microscope at 100X (Nikon AXR NSPARC).

### Protein expression and purification

The expression plasmid pTRIEX3neo containing the SATB1 full-length coding sequence or individual domains was transformed into the BL21 (DE3) bacterial cells. A single colony was inoculated in Luria-Bertani (LB) medium containing ampicillin and incubated at 37℃, 180 rpm for overnight. Next, 1% inoculum was added to 1 L auto-induction medium. The cells were grown for 5-6 h at 37°C, followed by 18℃ overnight at 180-200 rpm. The cells were harvested by centrifuging at 6000 X g for 20 min. The cell pellet was resuspended in Lysis Buffer (20 mM Tris-Cl pH 8.0, 300 mM NaCl, 0.2% Triton X-100, 10% Glycerol, 1 mM Lysozyme, 1 mM PMSF, 10 mM Imidazole and 1 mM DTT). The pellet was resuspended completely and incubated on ice for 15 min. Sonication of the sample was done with the following conditions: 60% amplitude - 2 sec ON/6 sec OFF – 10 min. The solution was centrifuged at 12000 rpm, 4℃, 30 min. The remaining supernatant is mixed with Ni-NTA beads (Takara) and incubated on the end-to-end rotator for 3 h, 8-10 rpm. The beads were washed 5-6 times with Wash Buffer (20 mM Tris-Cl pH 8.0, 300mM NaCl, 0.2% Triton X-100, 10% Glycerol, 20 mM Imidazole). The protein was eluted with Elution Buffer (20 mM Tris-Cl pH 8.0, 300 mM NaCl, 0.2% Triton X-100, 10% Glycerol, 100 mM Imidazole) 4 times and resolved on SDS-PAGE. The pooled fractions were dialyzed using 7-KDa cutoff dialysis bag in Dialysis Buffer I (20 mM Tris-Cl pH 8.0, 300 mM NaCl, 0.2% Triton X-100, 10% Glycerol) for 12-14 h. A buffer change to Dialysis Buffer II (20 mM Tris-Cl pH 8.0, 300 mM NaCl, 0.2% Triton X-100, 50% Glycerol) was done for 5-6 h. The protein was collected and stored at -80℃.

### In vitro droplet formation assay

The purified proteins were quantitated using BCA assay kit (Thermo Fisher Scientific) as per the manufacturers protocol. The droplet reactions were set up in 1.5 mL tubes as a 50 µL reaction with the following components: 1M HEPES pH 7-22 µl, TE pH 7.5 -12 µl, BSA-5.5 µl, 60% PEG-9 µl, 100 mM DTT-1.5 µl, DNA – 200 ng, Protein: variable. The solution, once it turned turbid, was imaged using a confocal microscope (Zeiss) with AF 488 and DIC channels. For droplet disruption experiments, 1,6-hexanediol (Sigma) was directly added to the reactions and incubated for 10, 15, and 30 min prior to imaging.

### Fluorescence recovery after photobleaching (FRAP) assay

The droplet formation assay described above was carried out. Upon visualization of droplets under fluorescence microscope, the points to be bleached were selected using the pointer tool in the FRAP settings of SP8 microscope (Leica). The following settings were used to image the selected points:

| Imaging parameters | pre-bleach | bleach | Post-bleach PB1 | Post bleach PB2 |
| --- | --- | --- | --- | --- |
| iterations | 1 |  | 15-20 | 15-20 |
| time/iteration | 0: 20: 640<br>min:sec:1/1000 | 0:7:400 | 0: 20: 640<br>min:sec:1/1000 | 0: 20: 640<br>min:sec:1/1000 |

### Databases and analyses for structure prediction

The amino acid sequences used for SATB1 and SMC1A were obtained from Uniprot database with accession numbers Q01826 and Q14683, respectively. The disorder prediction was carried out using PONDR (https://www.pondr.com/) with VL_X setting, as well as with IUPred (https://iupred2a.elte.hu/) with long-disorder option and PLAAC database (http://plaac.wi.mit.edu/) for prion-like similarity.

Alphafold2 using Google Colab2 notebook was used to predict de novo structures of SATB1 with no modifications to the default settings. The predicted structure of SMC1A was also obtained from the Alphafold database. The structures predicted were visualized in the PyMOL software.

### Site-directed mutagenesis

Point mutants in GFP-SATB1 were generated using site-directed mutagenesis procedure as described [93]. Briefly, amino acids that were identified to be mutated in various diseases were used to design oligonucleotides. One or multiple bases were mutated to gain the desired amino acid change using WT GFP-SATB1 in pEGFP-N1 mammalian expression vector. A two-step PCR reaction was set up with the conditions: denaturation 98°C for 5 min, cycled for 30 cycles with 98°C for 30 sec, 68°C for 8 min, and final extension 68°C for 10 min. The PCR products were run on 1% agarose gel. The desired DNA fragment was excised and gel extracted. The DNA was treated with DpnI at 37°C for 3 h to remove parental plasmids. The DNA was re-purified and transformed in DH5α ultra-competent bacteria. The colonies were screened by digestion with EcoRI. Positive clones were confirmed using Sanger sequencing. The oligonucleotide primers used for PCR are shown in Table S3.

### Cell culture, transfection and reporter assay

HEK293T cells were cultured in growth medium supplemented with 10% fetal bovine serum (Invitrogen) and maintained at 37°C in a humidified incubator with 5% CO₂. Transfections of mutant constructs were carried out using Lipofectamine 3000 (Invitrogen), following the manufacturer’s protocol. For the dual luciferase reporter assay, HEK293T cells were transfected with a reporter construct containing the *IL2* promoter cloned into the pGL3-basic vector. Co-transfections were performed with either empty vector, SATB1, SMC1A, or a combination of SMC1A and SATB1. To normalize for transfection efficiency, an equal amount of Renilla luciferase-expressing pTRK vector was included in each condition. After 48 h, cells were harvested and subjected to passive lysis, followed by the dual-luciferase assay (Promega) according to the manufacturer’s instructions. To evaluate the transcriptional activity of SATB1 mutants, wild-type or mutant SATB1 constructs were co-transfected with Il2-pGL3b, and reporter activity was assessed using the same assay protocol.

## Declarations

## Ethical approval

The research on animal models was conducted in accordance with the regulations of the Institutional Animal Ethics Committee and was approved by the same.

## Authors’ contributions

The project was conceived by SG, AM, and IP. The experimental design was contributed by SG, AM, IP, and GN. The experiments were performed by AM, IP, ID, and MT. VC and SV performed the proximity ligation assays. AM prepared all NGS libraries and data analysis, SS performed library quality checks, pooling and demultiplexing. The manuscript was written by AM and SG with inputs from IP, RL, and GN. Funding and resources were provided by SG, RL, and GN.

## Availability of data and materials

Sequencing data (ChIP-Seq and RNA-seq) is deposited in the Gene Expression Omnibus (GEO) and made publicly available upon peer-reviewed publication. Data and materials will be available from the lead contact upon reasonable request following publication.

## Acknowledgments

The authors acknowledge multiple funding agencies for supporting this work. The authors thank Dr Terumi Kohwi-Shigematsu (UCSF) for the gift of SATB1 floxed mice. We thank Dhrisaj Ray for help with protein purification. We thank the staff at the National Facility for Gene Function in Health and Disease at IISER Pune and Center for Integrative and Translational Research (CITRES), Shiv Nadar University for maintaining and providing the mice used in this study. We also thank the staff at the imaging facilities of IISER Pune and Shiv Nadar University.

