## Supplementary Information for "Satb1 integrates cohesin mediated genome organization and transcriptional regulation during T cell development"

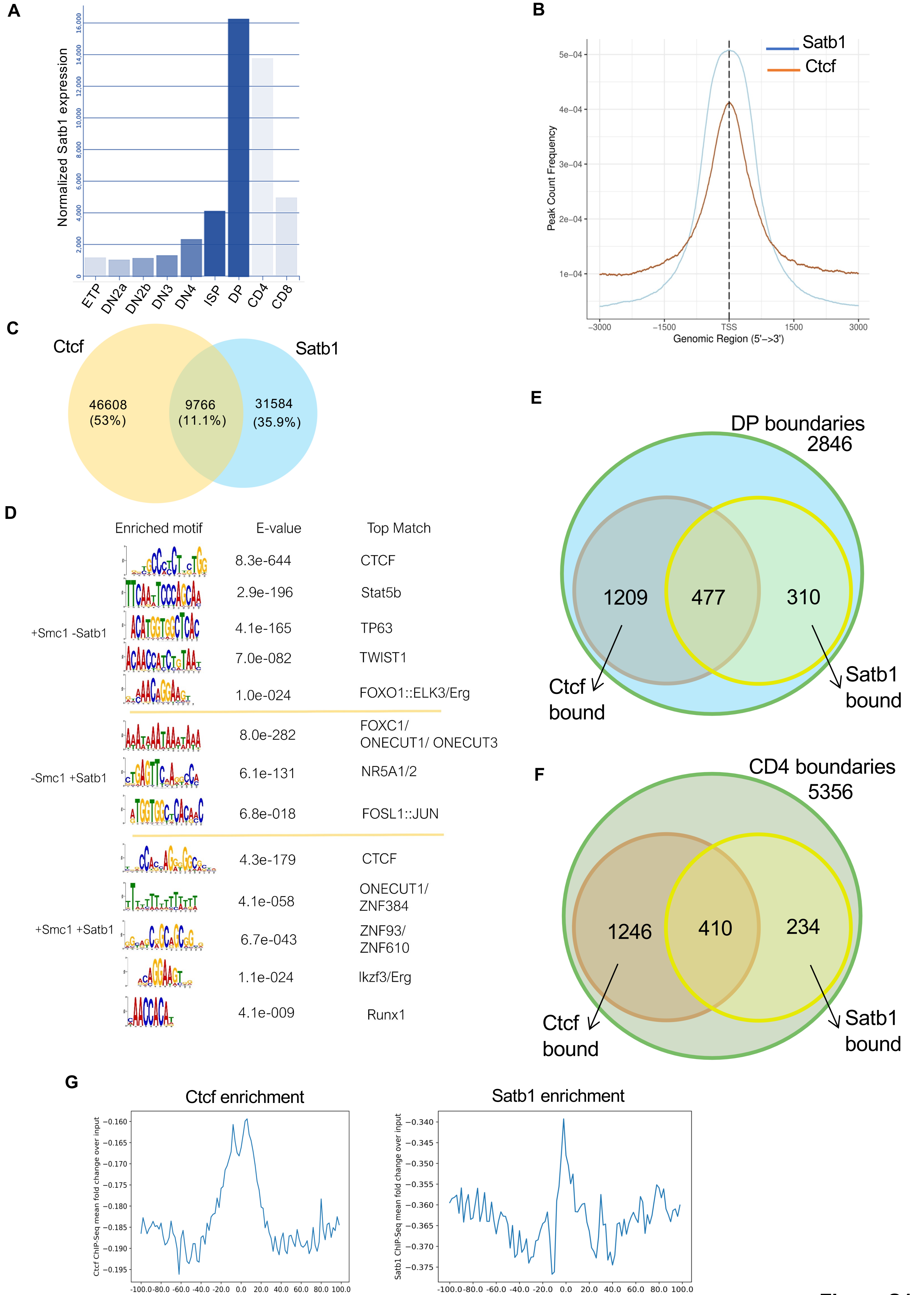
A
Normalized Satb1 expression
0
2,000
4,000
6,000
8,000
10,000
12,000
14,000
16,000
ETP
DN2a
DN2b
DN3
DN4
ISP
DP
CD4
CD8
B
Satb1
Ctcf
Peak Count Frequency
5e−04
4e−04
3e−04
2e−04
1e−04
−3000
−1500
TSS
1500
3000
Genomic Region (5'−>3')
C
Ctcf
Satb1
46608
(53%)
9766
(11.1%)
31584
(35.9%)
D
Enriched motif
E-value
Top Match
+Smc1 -Satb1
8.3e-644
CTCF
2.9e-196
Stat5b
4.1e-165
TP63
7.0e-082
TWIST1
1.0e-024
FOXO1::ELK3/Erg
-Smc1 +Satb1
8.0e-282
FOXC1/
ONECUT1/ ONECUT3
6.1e-131
NR5A1/2
6.8e-018
FOSL1::JUN
+Smc1 +Satb1
4.3e-179
CTCF
4.1e-058
ONECUT1/
ZNF384
6.7e-043
ZNF93/
ZNF610
1.1e-024
Ikzf3/Erg
4.1e-009
Runx1
E
DP boundaries
2846
1209
477
310
Ctcf
bound
Satb1
bound
F
CD4 boundaries
5356
1246
410
234
Ctcf
bound
Satb1
bound
G
Ctcf enrichment
Ctcf ChIP-Seq mean fold change over input
−0.160
−0.165
−0.170
−0.175
−0.180
−0.185
−0.190
−0.195
-100.0 -80.0 -60.0 -40.0 -20.0 0.0 20.0 40.0 60.0 80.0 100.0
Distance from boundary, kbp
Satb1 enrichment
Satb1 ChIP-Seq mean fold change over input
−0.340
−0.345
−0.350
−0.355
−0.360
−0.365
−0.370
−0.375
-100.0 -80.0 -60.0 -40.0 -20.0 0.0 20.0 40.0 60.0 80.0 100.0
Distance from boundary, kbp

**Figure S1**

**Figure S1. Satb1 shares genomic occupancy with Ctcf in DP thymocytes. A**. Normalized Satb1 expression profile across thymic developmental stages using publicly available datasets [95]. **B**. ChIP-seq peaks of Satb1 and Ctcf overlayed as average profiles around the TSS. Distance from TSS on either side is ±3 kb. **C**. Number of common and unique binding sites of Satb1 and Ctcf in DP thymocytes. **D**. Motif enrichment analysis of ChIP-seq peaks; Smc1a excluding Satb1, Satb1 without Smc1a, and the common peaks between the two proteins, respectively. E. TAD boundaries were called for DP thymocyte HiC sample. The venn diagram shown indicates the number of boundaries co-occupied by Satb1 and Ctcf in DP cells uniquely, as well as depicting their common boundaries. **F**. Similar to **E**, boundaries appearing in CD4 (CD69+) HiC dataset were intersected with Ctcf and Satb1 binding sites to obtain their overlap at CD4 thymocyte boundary regions. **G**. Average distance of Ctcf and Satb1 ChIP binding from DP specific insulation boundaries, respectively. Fold-change over input for the factors was used to calculate the distance from boundary center with a flanking region of 100 kb on either side of the boundary is shown.

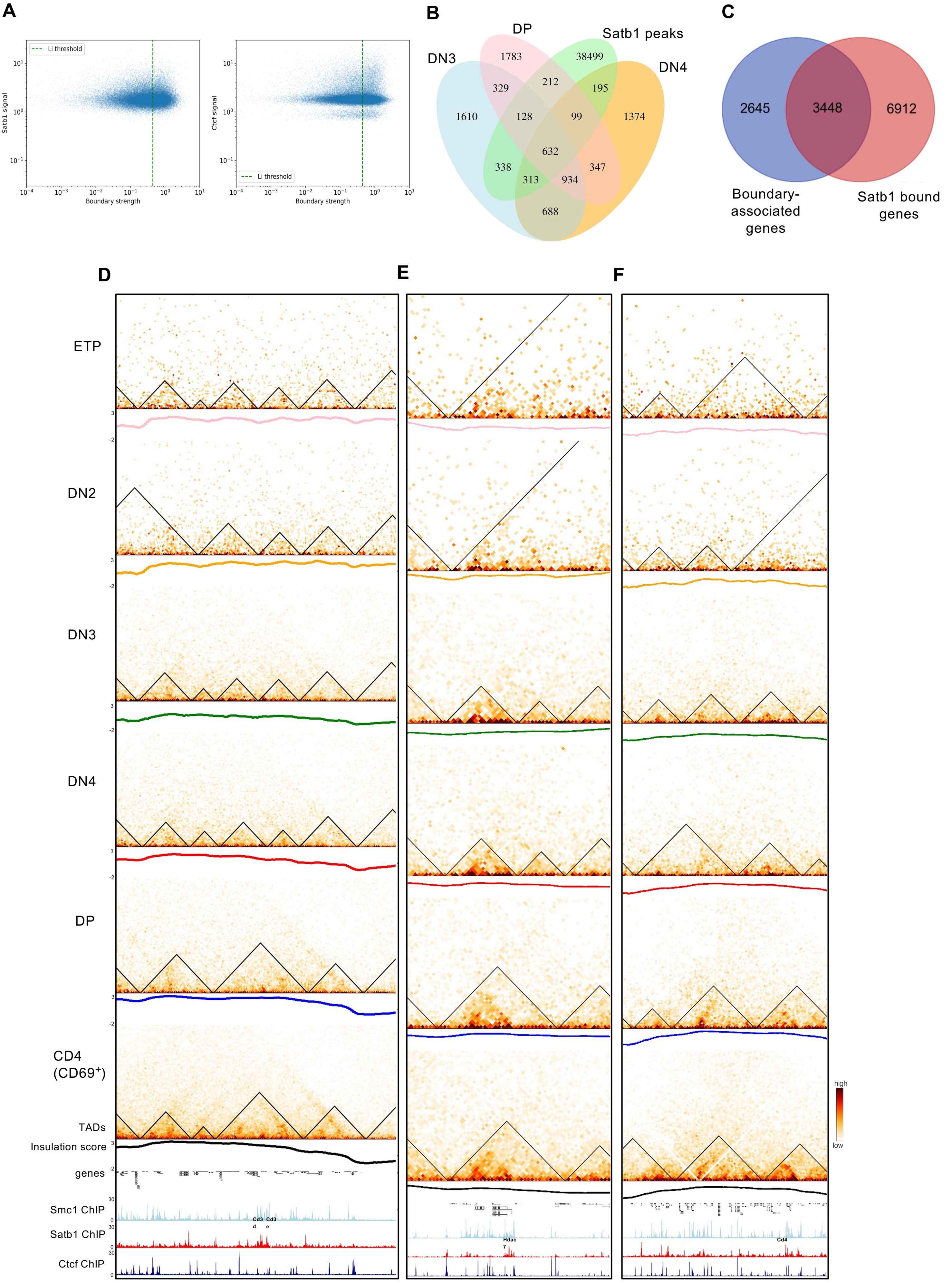
A
Li threshold
Satb1 signal
Ctcf signal
Boundary strength
B
DP
Satb1 peaks
DN3
DN4
1783
38499
329
212
195
1610
128
99
1374
632
338
347
313
934
688
C
2645
3448
6912
Boundary-associated genes
Satb1 bound genes
D
E
F
ETP
DN2
DN3
DN4
DP
CD4 (CD69⁺)
TADs
Insulation score
genes
Smc1 ChIP
Satb1 ChIP
Ctcf ChIP
Cd3d
Cd3e
Hdac7
Cd4
high
low

Figure S2

**Figure S2. TAD boundaries are marked by Satb1 along with Ctcf and Cohesin. A**. Boundary strength of DP HiC data was plotted along normalized ChIP-seq signal of Satb1 (top) and Ctcf (bottom) in DP thymocytes. Cooltools [53] were used to calculate insulation score and boundary strength threshold. **B**. Overlap of TAD boundaries specific to DN3, DN4 and DP thymocytes with Satb1 ChIP-seq peaks where at least one anchor is associated with Satb1. C. Venn diagram showing intersection of Satb1 bound genes and boundary associated genes in DP thymocytes. Genomic interactions were shown at the *Cd3* (**D**), *Hdac7* (**E**) and *Cd4* (**F**) loci in various T-cell developmental stages. Below each HiC contact matrix, the stage specific insulation score is plotted. The panels below depict genome browser view of Ctcf, Cohesin (Smc1a) and Satb1 occupancies in DP thymocytes.

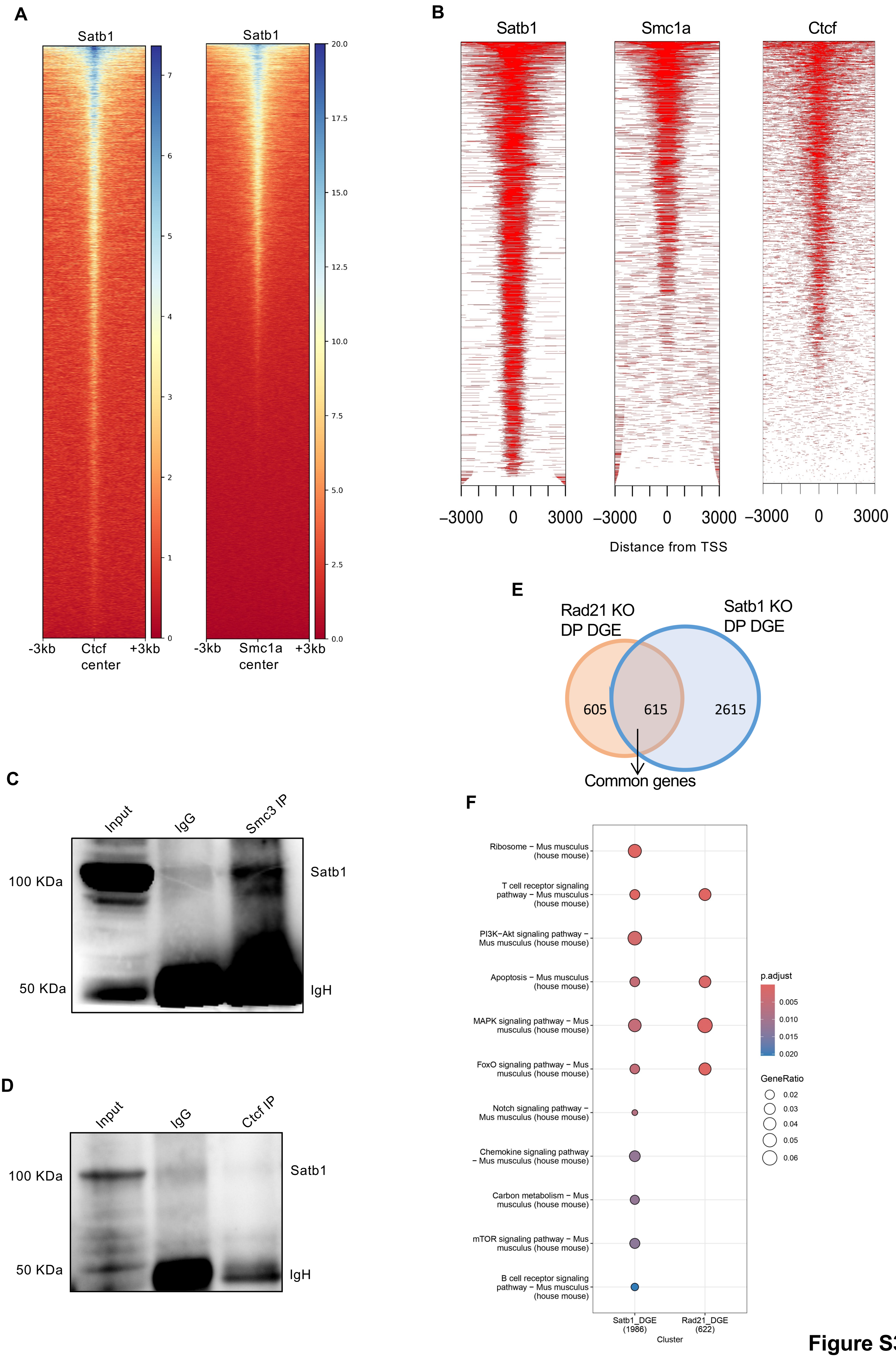
A
Satb1
Satb1
-3kb
Ctcf center
+3kb
-3kb
Smc1a center
+3kb
B
Satb1
Smc1a
Ctcf
−3000
0
3000
Distance from TSS
C
Input
IgG
Smc3 IP
100 KDa
50 KDa
Satb1
IgH
D
Input
IgG
Ctcf IP
100 KDa
50 KDa
Satb1
IgH
E
Rad21 KO DP DGE
Satb1 KO DP DGE
605
615
2615
Common genes
F
Ribosome − Mus musculus (house mouse)
T cell receptor signaling pathway − Mus musculus (house mouse)
PI3K−Akt signaling pathway − Mus musculus (house mouse)
Apoptosis − Mus musculus (house mouse)
MAPK signaling pathway − Mus musculus (house mouse)
FoxO signaling pathway − Mus musculus (house mouse)
Notch signaling pathway − Mus musculus (house mouse)
Chemokine signaling pathway − Mus musculus (house mouse)
Carbon metabolism − Mus musculus (house mouse)
mTOR signaling pathway − Mus musculus (house mouse)
B cell receptor signaling pathway − Mus musculus (house mouse)
Satb1_DGE (1986)
Rad21_DGE (622)
Cluster
p.adjust
0.005
0.010
0.015
0.020
GeneRatio
0.02
0.03
0.04
0.05
0.06

**Figure S3**

**Figure S3. Satb1 and cohesin binding profiles share multiple genomic features in DP thymocytes**. **A**. Heatmap of normalized ChIP-seq profile of Satb1 relative to Ctcf (left) and Smc1a (right) binding sites in DP thymocytes. **B**. Occupancy of Satb1, Smc1a and Ctcf relative to TSS are shown as tag-density heatmaps. **C**. Satb1 was found to co-immunoprecipitate with Smc3, another core component of the cohesin complex. Here, Smc3 was immunoprecipitated, and Satb1 was detected by immunoblotting. **D**. Ctcf IP was performed in DP thymocytes followed by Satb1 WB. **E**. Publicly available RNA-seq of Rad21 KO in DP thymocytes was analysed and DEGs obtained were compared with genes significantly dysregulated in Satb1 KO DPs. **F**. Significantly enriched KEGG pathways for Rad21KO/WT and Satb1 KO/WT were clustered and compared. Top 10 pathways with padj <0.05 are depicted.

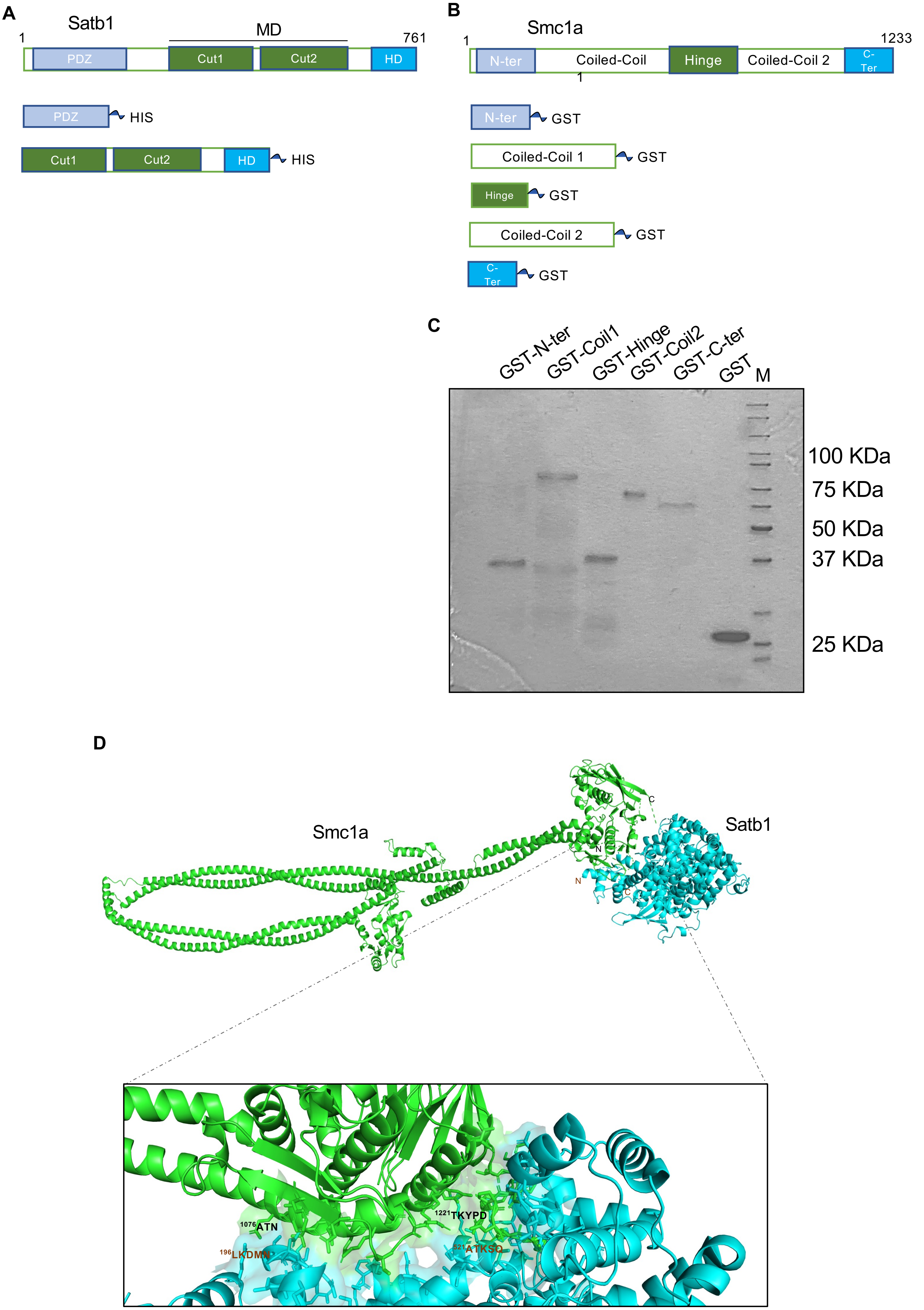
A
Satb1
1
MD
761
PDZ
Cut1
Cut2
HD
PDZ
HIS
Cut1
Cut2
HD
HIS
B
Smc1a
1
1233
N-ter
Coiled-Coil
1
Hinge
Coiled-Coil 2
C-Ter
N-ter
GST
Coiled-Coil 1
GST
Hinge
GST
Coiled-Coil 2
GST
C-Ter
GST
C
GST-N-ter
GST-Coil1
GST-Hinge
GST-Coil2
GST-C-ter
GST
M
100 KDa
75 KDa
50 KDa
37 KDa
25 KDa
D
Smc1a
Satb1
C
N
N
C
1076ATN
1221TKYPD
196LKDMN
521ATKSQ

**Figure S4**

**Figure S4. Dissection of interaction domains of Satb1 and Smc1a. A.** Schematic of full length SATB1 and major domains of Satb1 which were cloned as poly-histidine (HIS) tagged ORFs in pTRIEX-3 Neo. **B.** Schematic representation of full-length and the functional domains of Smc1a. The individual domains, as depicted, were cloned into pGEX4T1 vector to tag them with glutathione S transferase (GST) gene present in the vector. **C.** The individual domains of Smc1a were purified as GST-fusion proteins using glutathione beads. Panel depicts denaturing polyacrylamide gel (SDS-PAGE) stained with Coomassie blue. Lane 'M' denotes the molecular weight marker. **D**. Predicted 3D structure of heterodimeric interaction of Smc1a (green) and Satb1 (blue) using Alphafold2. The structure was visualized in PyMol. Lower panel shows zoomed in region showing the predicted interaction residues of Smc1a and Satb1. The primary interaction residues are labelled; Smc1a residues-black, Satb1 residues-brown.

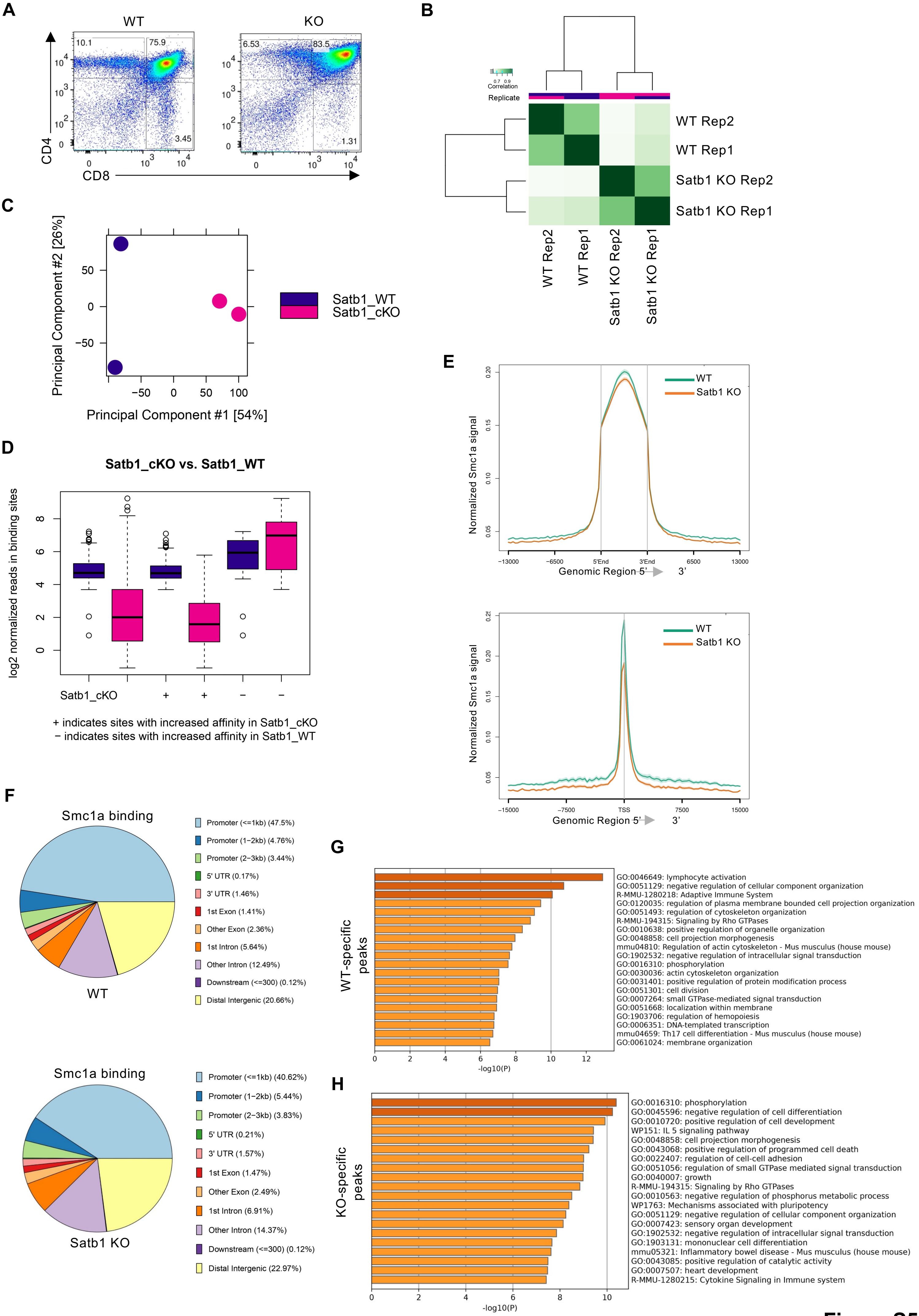
A
WT
KO
CD4
CD8
10.1
75.9
3.45
6.53
83.5
1.31
B
Correlation
Replicate
WT Rep2
WT Rep1
Satb1 KO Rep2
Satb1 KO Rep1
C
Principal Component #2 [26%]
Principal Component #1 [54%]
Satb1_WT
Satb1_cKO
D
Satb1_cKO vs. Satb1_WT
log2 normalized reads in binding sites
Satb1_cKO
+ indicates sites with increased affinity in Satb1_cKO
− indicates sites with increased affinity in Satb1_WT
E
Normalized Smc1a signal
WT
Satb1 KO
Genomic Region 5' 3'
5'End
3'End
TSS
F
Smc1a binding
WT
Promoter (<=1kb) (47.5%)
Promoter (1−2kb) (4.76%)
Promoter (2−3kb) (3.44%)
5' UTR (0.17%)
3' UTR (1.46%)
1st Exon (1.41%)
Other Exon (2.36%)
1st Intron (5.64%)
Other Intron (12.49%)
Downstream (<=300) (0.12%)
Distal Intergenic (20.66%)
Smc1a binding
Satb1 KO
Promoter (<=1kb) (40.62%)
Promoter (1−2kb) (5.44%)
Promoter (2−3kb) (3.83%)
5' UTR (0.21%)
3' UTR (1.57%)
1st Exon (1.47%)
Other Exon (2.49%)
1st Intron (6.91%)
Other Intron (14.37%)
Downstream (<=300) (0.12%)
Distal Intergenic (22.97%)
G
WT-specific peaks
GO:0046649: lymphocyte activation
GO:0051129: negative regulation of cellular component organization
R-MMU-1280218: Adaptive Immune System
GO:0120035: regulation of plasma membrane bounded cell projection organization
GO:0051493: regulation of cytoskeleton organization
R-MMU-194315: Signaling by Rho GTPases
GO:0010638: positive regulation of organelle organization
GO:0048858: cell projection morphogenesis
mmu04810: Regulation of actin cytoskeleton - Mus musculus (house mouse)
GO:1902532: negative regulation of intracellular signal transduction
GO:0016310: phosphorylation
GO:0030036: actin cytoskeleton organization
GO:0031401: positive regulation of protein modification process
GO:0051301: cell division
GO:0007264: small GTPase-mediated signal transduction
GO:0051668: localization within membrane
GO:1903706: regulation of hemopoiesis
GO:0006351: DNA-templated transcription
mmu04659: Th17 cell differentiation - Mus musculus (house mouse)
GO:0061024: membrane organization
-log10(P)
H
KO-specific peaks
GO:0016310: phosphorylation
GO:0045596: negative regulation of cell differentiation
GO:0010720: positive regulation of cell development
WP151: IL 5 signaling pathway
GO:0048858: cell projection morphogenesis
GO:0043068: positive regulation of programmed cell death
GO:0022407: regulation of cell-cell adhesion
GO:0051056: regulation of small GTPase mediated signal transduction
GO:0040007: growth
R-MMU-194315: Signaling by Rho GTPases
GO:0010563: negative regulation of phosphorus metabolic process
WP1763: Mechanisms associated with pluripotency
GO:0051129: negative regulation of cellular component organization
GO:0007423: sensory organ development
GO:1902532: negative regulation of intracellular signal transduction
GO:1903131: mononuclear cell differentiation
mmu05321: Inflammatory bowel disease - Mus musculus (house mouse)
GO:0043085: positive regulation of catalytic activity
GO:0007507: heart development
R-MMU-1280215: Cytokine Signaling in Immune system
-log10(P)

**Figure S5**

**Figure S5**. **Genomic occupancy of Smc1a exhibits reduced preference at promoters upon Satb1 deletion**. **A.** Flow cytometry analysis showing the effects of Satb1 depletion on thymocyte populations. Representative plot from 3 independent experiments. **B**. Distance correlation heatmap of Smc1a Cut&Run in WT and Satb1 KO samples was shown. **C**. Principal component analysis depicting the global differences in Smc1a binding in WT vs KO. **D**. The increased or decreased affinity of Smc1a binding in WT and Satb1 KO DP thymocytes is shown as boxplot. **E**. Normalized Smc1a signal at Satb1 binding sites and TSS regions, respectively, are shown as average line plots. **F**. Genomic annotations of Smc1a binding in WT (top) and Satb1 KO (bottom) are shown as pie-charts. Gene ontology analysis shows top biological processes associated with Smc1a peaks in WT (**G**) and Satb1 KO (**H**) conditions.

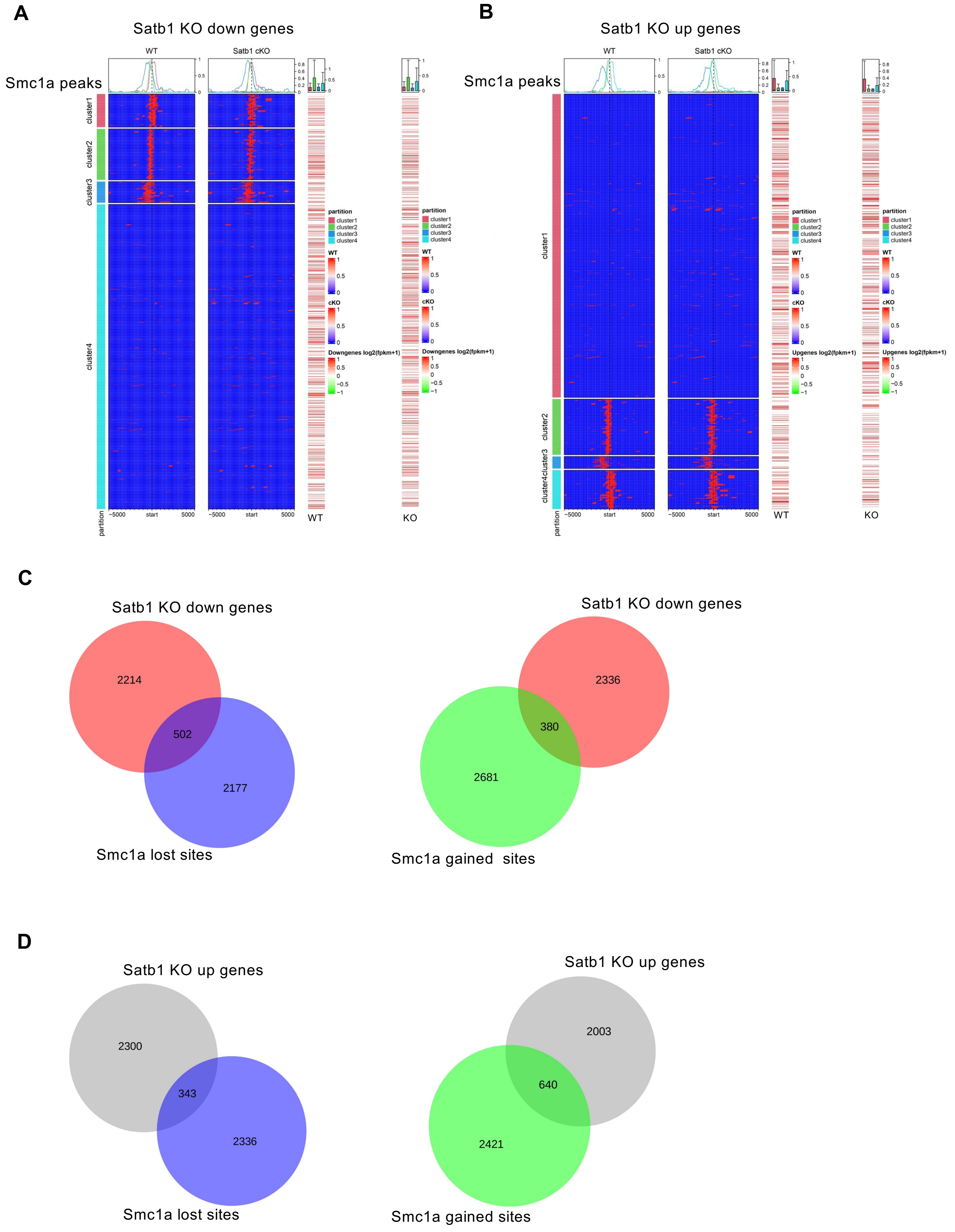
A
Satb1 KO down genes
Smc1a peaks
WT
Satb1 cKO
cluster1
cluster2
cluster3
cluster4
partition
−5000
start
5000
WT
KO
partition
cluster1
cluster2
cluster3
cluster4
WT
cKO
Downgenes log2(fpkm+1)
B
Satb1 KO up genes
Smc1a peaks
WT
Satb1 cKO
Upgenes log2(fpkm+1)
C
Satb1 KO down genes
2214
502
2177
Smc1a lost sites
Satb1 KO down genes
2336
380
2681
Smc1a gained sites
D
Satb1 KO up genes
2300
343
2336
Smc1a lost sites
Satb1 KO up genes
2003
640
2421
Smc1a gained sites

**Figure S6**

**Figure S6. Correlation of Smc1a binding and Satb1 mediated gene expression in DP thymocytes. A.** k-means clustering of Smc1a peaks enriched in WT and Satb1 KO datasets and clustered alongside transcript levels of downregulated genes in Satb1 KO DPs, both in WT and KO samples. **B**. Clustering similar to **A**, using normalized transcript expression of upregulated genes upon Satb1 KO. RPKM values are depicted as heatmap of RNA-seq data. **C**. Venn diagrams depicting the commonality between genes downregulated and gained or lost Smc1a peaks upon Satb1 KO compared to WT. **D**. Similar comparison was done with upregulated genes upon Satb1 KO.

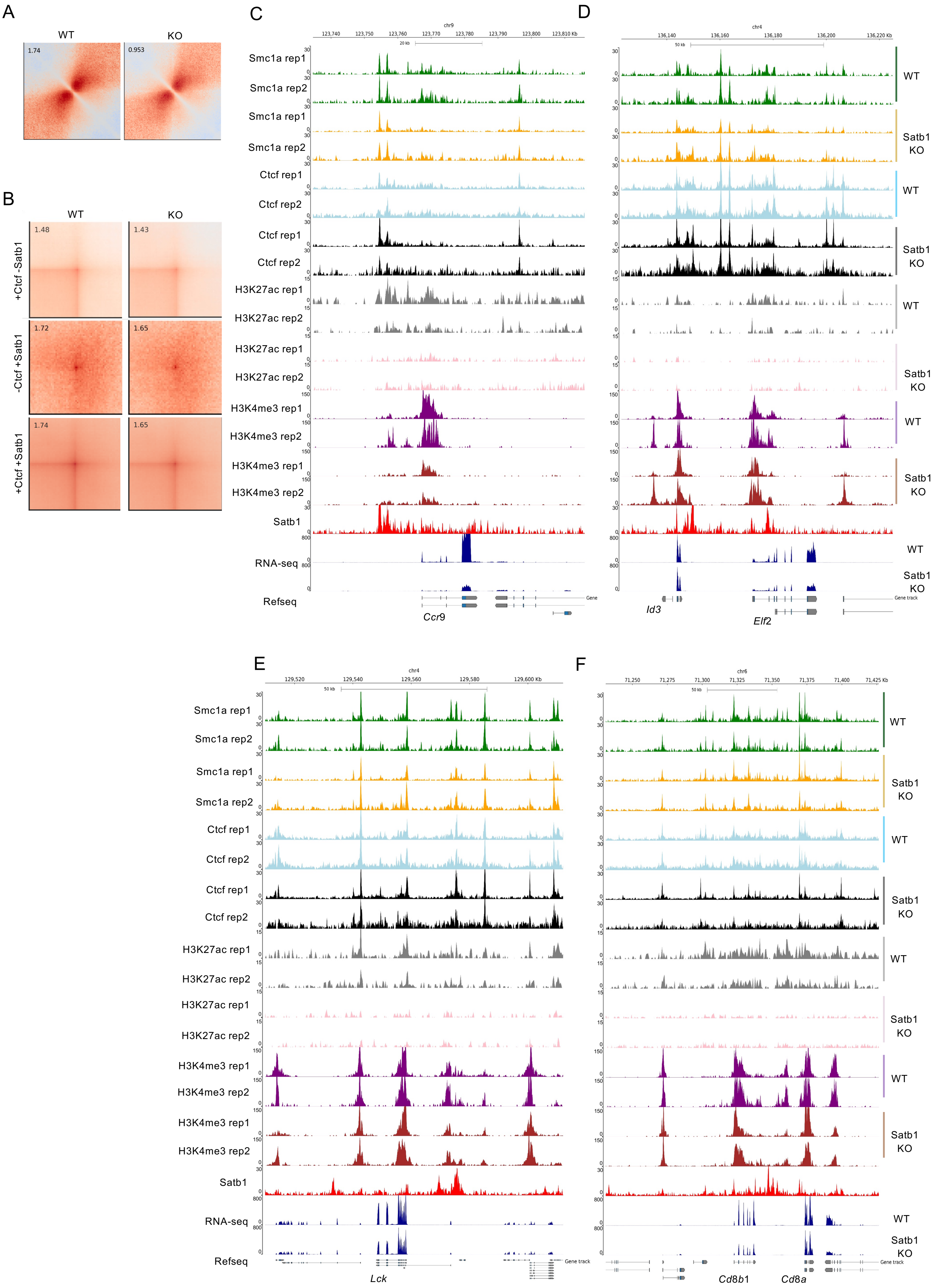
A
WT
KO
1.74
0.953
B
WT
KO
+Ctcf -Satb1
-Ctcf +Satb1
+Ctcf +Satb1
1.48
1.43
1.72
1.65
1.74
1.65
C
chr9
Smc1a rep1
Smc1a rep2
Smc1a rep1
Smc1a rep2
Ctcf rep1
Ctcf rep2
Ctcf rep1
Ctcf rep2
H3K27ac rep1
H3K27ac rep2
H3K27ac rep1
H3K27ac rep2
H3K4me3 rep1
H3K4me3 rep2
H3K4me3 rep1
H3K4me3 rep2
Satb1
RNA-seq
Refseq
Ccr9
D
chr4
WT
Satb1 KO
WT
Satb1 KO
WT
Satb1 KO
WT
Satb1 KO
WT
Satb1 KO
Id3
Elf2
E
chr4
Smc1a rep1
Smc1a rep2
Smc1a rep1
Smc1a rep2
Ctcf rep1
Ctcf rep2
Ctcf rep1
Ctcf rep2
H3K27ac rep1
H3K27ac rep2
H3K27ac rep1
H3K27ac rep2
H3K4me3 rep1
H3K4me3 rep2
H3K4me3 rep1
H3K4me3 rep2
Satb1
RNA-seq
Refseq
Lck
F
chr6
WT
Satb1 KO
WT
Satb1 KO
WT
Satb1 KO
WT
Satb1 KO
WT
Satb1 KO
Cd8b1
Cd8a

**Figure S7**

**Figure S7. Smc1a occupancy at TSS of TCR specific genes decreases upon Satb1 KO**. **A**. Local pileup analysis shows genomic interactions at Satb1 only sites in WT and Satb1 KO conditions. A 600 kb padding region was given for the estimation. **B**. Genomic interactions at Satb1, Ctcf and Satb1+Ctcf sites in WT and Satb1 KO conditions are shown by pileup analysis. Genome browser view of Smc1a and Ctcf occupancy along with histone modifications H3K27ac and H3K4me3, in WT and Satb1 KO conditions are shown at *Ccr9* (**C**), *Elf2* (**D**), *Lck* (**E**) and *Cd8* (**F**) loci. Normalized transcript expression is also shown for these genes in WT and Satb1 KO conditions (bottom).

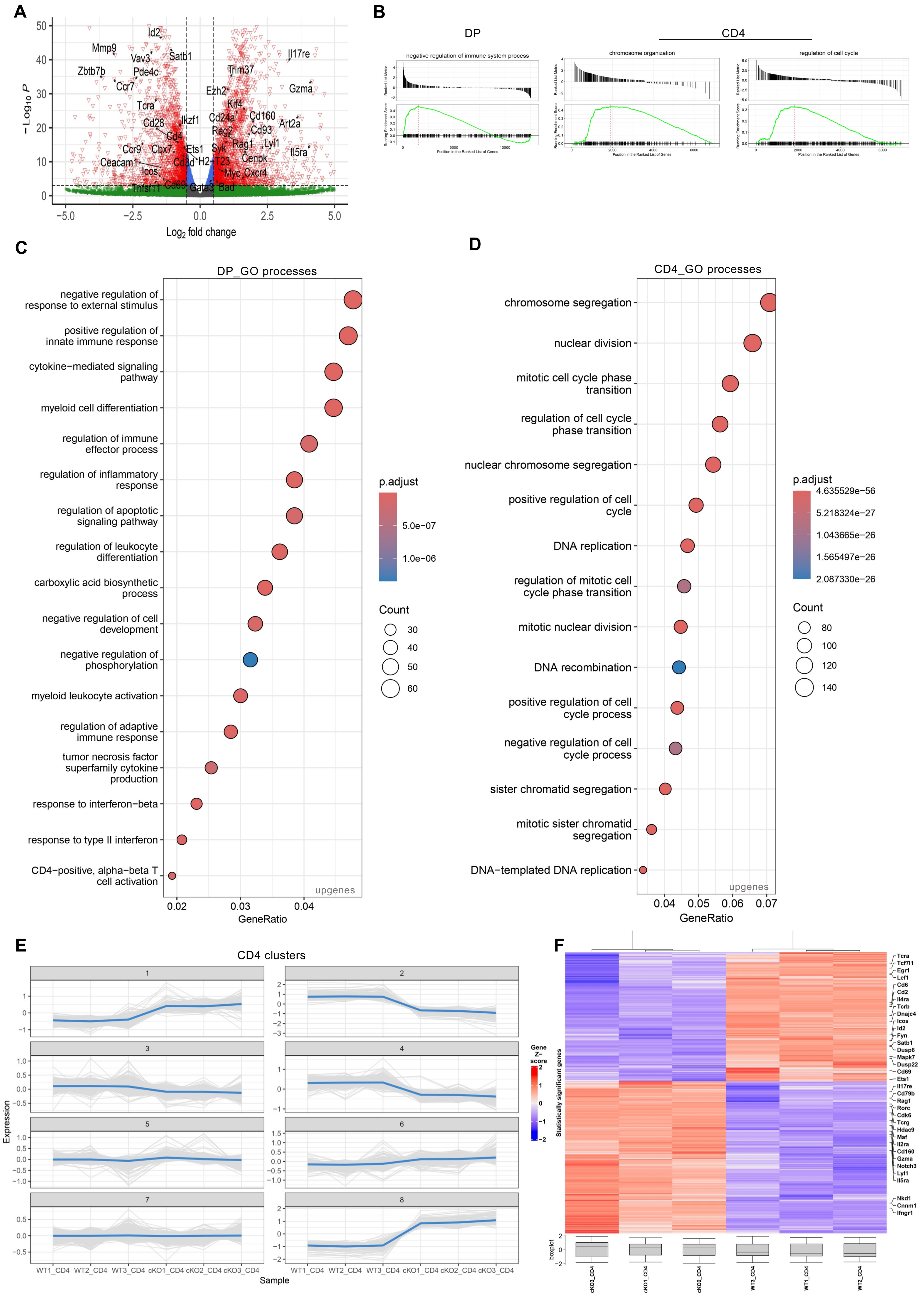
A
B
DP
CD4
negative regulation of immune system process
chromosome organization
regulation of cell cycle
Position in the Ranked List of Genes
C
DP_GO processes
negative regulation of response to external stimulus
positive regulation of innate immune response
cytokine−mediated signaling pathway
myeloid cell differentiation
regulation of immune effector process
regulation of inflammatory response
regulation of apoptotic signaling pathway
regulation of leukocyte differentiation
carboxylic acid biosynthetic process
negative regulation of cell development
negative regulation of phosphorylation
myeloid leukocyte activation
regulation of adaptive immune response
tumor necrosis factor superfamily cytokine production
response to interferon−beta
response to type II interferon
CD4−positive, alpha−beta T cell activation
upgenes
GeneRatio
p.adjust
5.0e−07
1.0e−06
Count
30
40
50
60
D
CD4_GO processes
chromosome segregation
nuclear division
mitotic cell cycle phase transition
regulation of cell cycle phase transition
nuclear chromosome segregation
positive regulation of cell cycle
DNA replication
regulation of mitotic cell cycle phase transition
mitotic nuclear division
DNA recombination
positive regulation of cell cycle process
negative regulation of cell cycle process
sister chromatid segregation
mitotic sister chromatid segregation
DNA−templated DNA replication
upgenes
GeneRatio
p.adjust
4.635529e−56
5.218324e−27
1.043665e−26
1.565497e−26
2.087330e−26
Count
80
100
120
140
E
CD4 clusters
Expression
Sample
WT1_CD4 WT2_CD4 WT3_CD4 cKO1_CD4 cKO2_CD4 cKO3_CD4
F
Gene Z−score
Statistically significant genes
boxplot
cKO3_CD4
cKO1_CD4
cKO2_CD4
WT3_CD4
WT1_CD4
WT2_CD4

**Figure S8**

**Figure S8. Effect of Satb1 depletion in CD4 thymocytes. A.** Examples of positively enriched processes in DP and CD4 upon Satb1 KO using functional GSEA. **B**. Volcano plot showing up- and downregulated genes in CD4 thymocytes upon Satb1 depletion. n=3 was used for analysis. **C**. Top GO terms enriched in DP (left) and **D**. CD4 (right) upregulated genes upon Satb1 knockout. **E**. k-means clustering of CD4 datasets in WT and Satb1 KO condition to depict groups of transcriptional changes in the two conditions (n=3). **F**. Heatmap showing the top genes dysregulated in CD4 thymocytes in Satb1 KO compared to WT cells. The overall transcripts change comparison is shown at the bottom as boxplots.

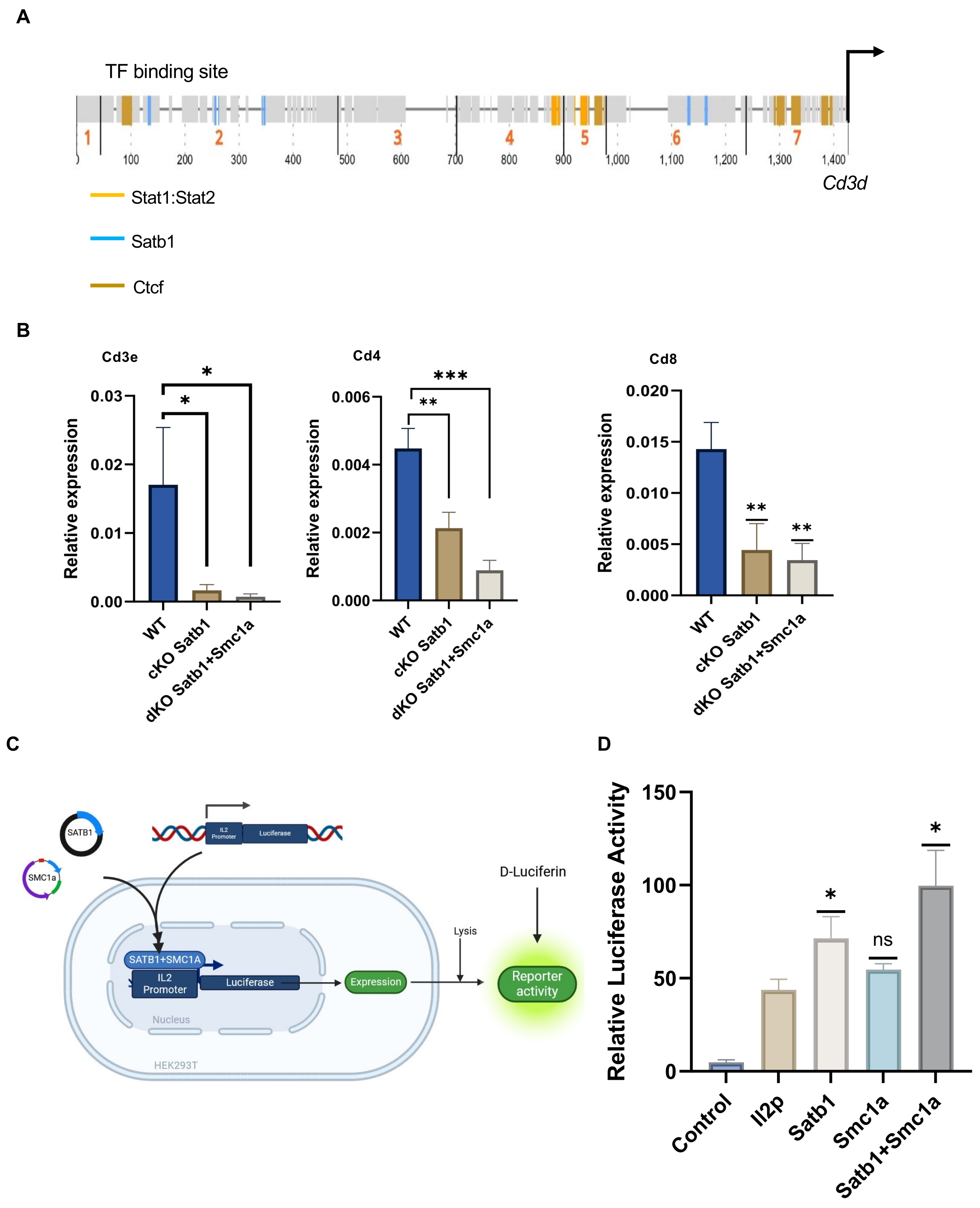
A
TF binding site
1
2
3
4
5
6
7
100
200
300
400
500
600
700
800
900
1,000
1,100
1,200
1,300
1,400
Cd3d
Stat1:Stat2
Satb1
Ctcf
B
Cd3e
Cd4
Cd8
Relative expression
0.03
0.02
0.01
0.00
0.006
0.004
0.002
0.000
0.020
0.015
0.010
0.005
0.000
*
**
***
WT
cKO Satb1
dKO Satb1+Smc1a
C
SATB1
SMC1a
IL2 Promoter
Luciferase
SATB1+SMC1A
Expression
Nucleus
HEK293T
Lysis
D-Luciferin
Reporter activity
D
Relative Luciferase Activity
150
100
50
0
ns
Control
Il2p
Satb1
Smc1a
Satb1+Smc1a

Figure S9

**Figure S9. Satb1 mediated transcriptional regulation is augmented by cohesin at T lineage specific genes**. **A.** Known-motif identification at *Cd3d* promoter using ConTra v3 tool [96]. **B**. RT-qPCR expression profiling of three target genes, enriched for both Smc1a and Satb1 binding, in WT, Satb1 KO and Satb1+Smc1a KO (dKO) conditions. All the three genes display a reduced expression in Satb1 KO, expression of which are further decreased, in the dKO condition. Expression of 18s rRNA was used to normalize gene expression, n=3. **C.** Schematic of luciferase assay utilized to assess the collaborative role of Satb1 and Smc1a in vitro. **D.** Relative luciferase activity by *Il2* promoter under different transfection combinations in HEK293T. Measurements of renilla (pTRK) activity was used to normalize the luciferase activity, n=4. Statistical significance was calculated using ANOVA in Graphpad v10. * $p<0.05$, ** $p<0.01$, *** $p<0.001$.

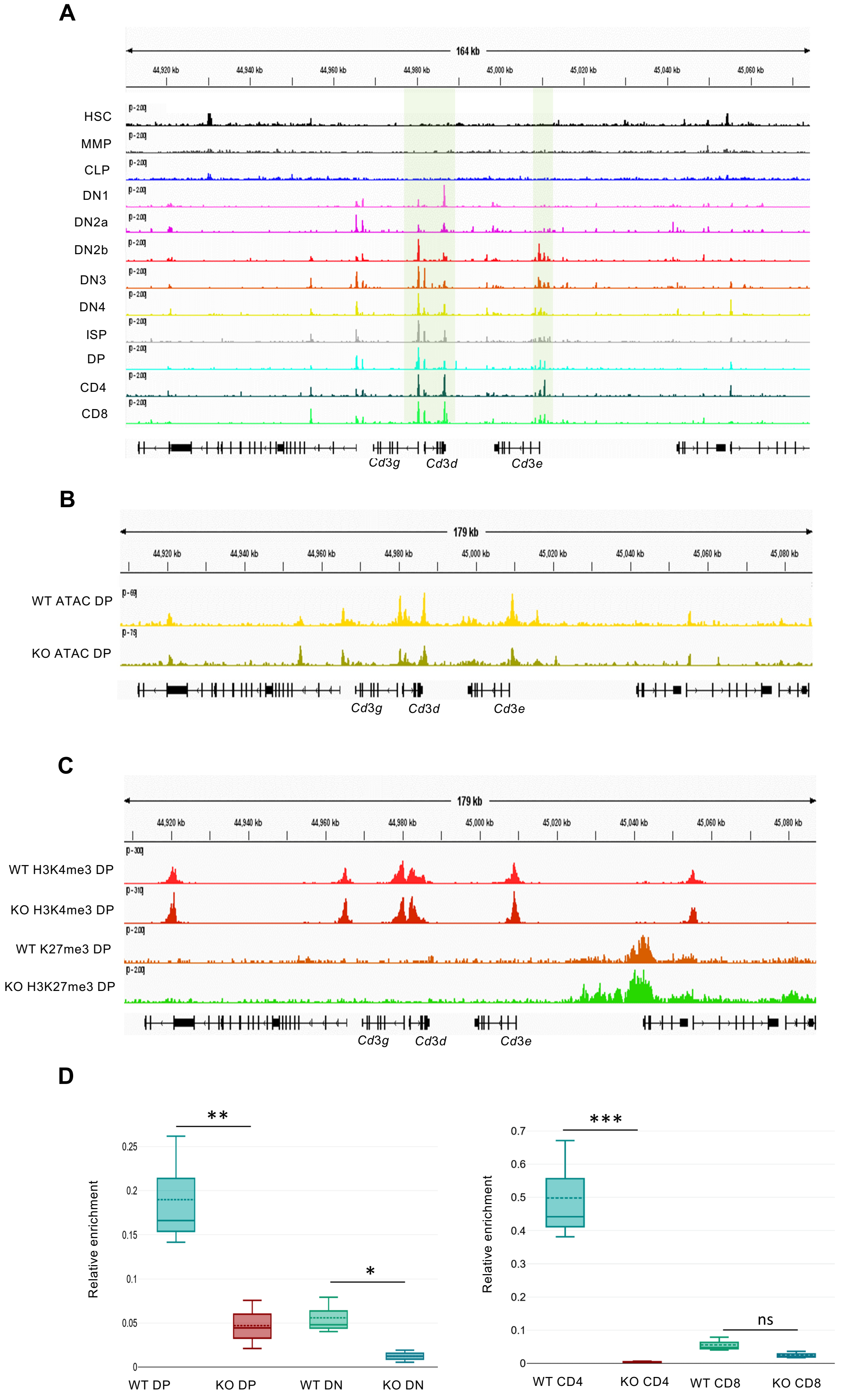
A
164 kb
HSC
MMP
CLP
DN1
DN2a
DN2b
DN3
DN4
ISP
DP
CD4
CD8
Cd3g
Cd3d
Cd3e
B
179 kb
WT ATAC DP
KO ATAC DP
Cd3g
Cd3d
Cd3e
C
179 kb
WT H3K4me3 DP
KO H3K4me3 DP
WT K27me3 DP
KO H3K27me3 DP
Cd3g
Cd3d
Cd3e
D
Relative enrichment
**
*
WT DP
KO DP
WT DN
KO DN
Relative enrichment
***
ns
WT CD4
KO CD4
WT CD8
KO CD8

Figure S10

**Figure S10. Satb1 is essential for chromatin accessibility at the *Cd3* locus during T cell development**. **A**. ATACseq analysis of publicly available data of various pre-T, pro-T and committed T lineage cells, (as indicated) at the *Cd3* locus is shown as IGV tracks. **B**. IGV tracks at the *Cd3* locus shows changes in chromatin accessibility in WT and Satb1 KO samples [39]. **C**. IGV tracks with histone modifications (as indicated) shows epigenetic modulation at the Cd3 locus because of Satb1 depletion in DP thymocytes. **D**. Assay for transposase-accessible chromatin (ATAC) was performed in WT and Satb1 KO sorted thymocytes, followed by quantitative RT-PCR. Chromatin accessibility at *Cd3e* promoter is shown for DN, DP CD8 and CD4 populations for WT and Satb1 KO cells. Actin promoter was used for normalization. Two independent replicates were used for statistical significance using ANOVA. * $p<0.05$, ** $p<0.01$, *** $p<0.001$.

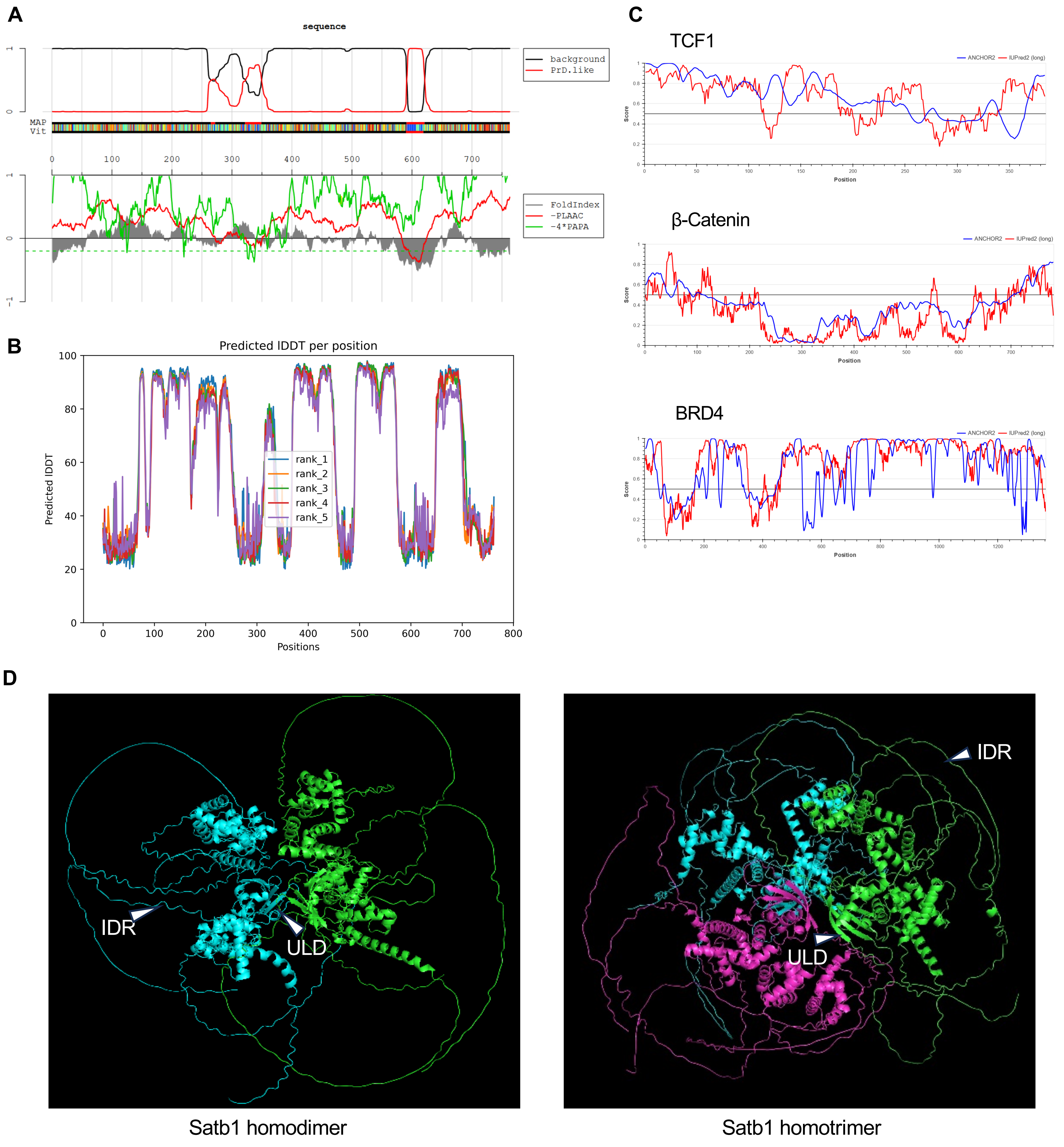

A
sequence
background
PrD.like
MAP
Vit
FoldIndex
-PLAAC
-4*PAPA
B
Predicted lDDT per position
Predicted lDDT
Positions
rank_1
rank_2
rank_3
rank_4
rank_5
C
TCF1
β-Catenin
BRD4
ANCHOR2
IUPred2 (long)
Score
Position
D
IDR
ULD
Satb1 homodimer
Satb1 homotrimer


Figure S11

**Figure S11. In silico analysis of Satb1 primary sequence reveals low-complexity and intrinsically disordered regions**. **A** PLAAC analysis (http://plaac.wi.mit.edu/) of Satb1 amino acid sequence is shown. Similarity to prion-like region is indicated (red). **B**. In silico analysis (using Alphafold) [97] of SATB1 sequence showing confidence scores for accurate structure prediction. A higher IDDT score depicts more structured regions, and the lower scores depicts IDR-like regions in the amino acid sequence. **C**. Similar disorder prediction analyses on other factors known to phase separate in vitro using IUPRED webtool [94]. **D**. De novo structure prediction [97] of monomeric SATB1 protein using its amino acid sequence. A similar structure prediction analysis was done for trimeric SATB1 (right), showing that the predicted IDRs could interact with other IDRs as well as functional domains of SATB1. ULD domain and IDRs are indicated with arrowheads.

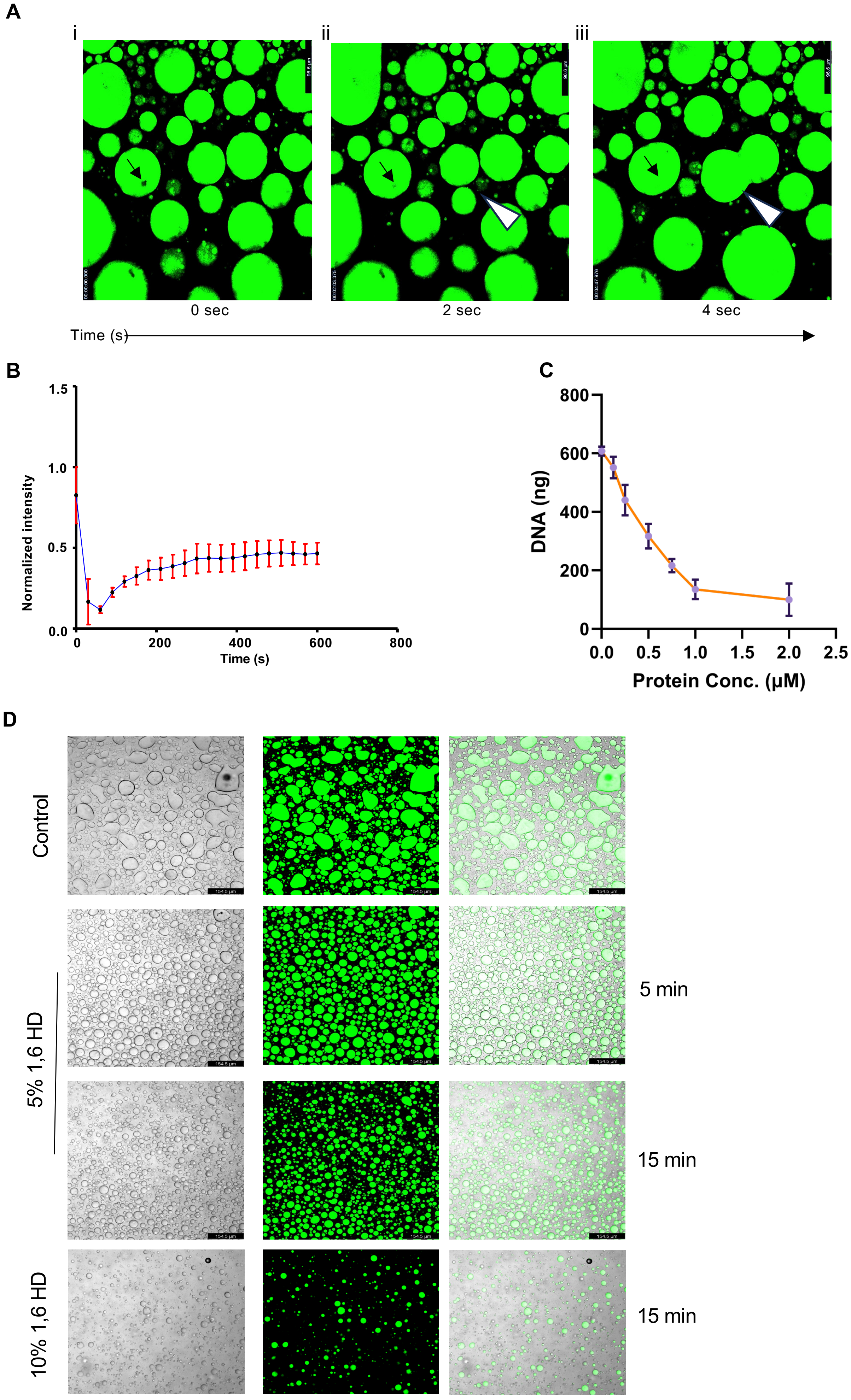
A
i
ii
iii
0 sec
2 sec
4 sec
Time (s)
B
Normalized intensity
Time (s)
C
DNA (ng)
Protein Conc. (μM)
D
Control
5% 1,6 HD
10% 1,6 HD
5 min
15 min
15 min

Figure S12

**Figure S12. Satb1 displays biophysical properties of liquid-liquid phase separation (LLPS) in vitro. A.** Droplet formation assay for Satb1 was performed in vitro using 0.5μM protein in the presence of PEG. Fluorescence recovery after photobleaching (FRAP) was performed followed by continuous imaging upto 20 min. The bleach point is marked inside the droplet in (i). (ii) shows immediate recovery at the bleach site. (iii) At the bottom, marked by an arrowhead, two droplets show fusion or merging behavior to coalesce into a bigger droplet. **B.** Quantification of the recovery in the fluorescence intensity at the fluorescence depleted point, shown in (i), as a function of time. **C.** In vitro droplets were centrifuged. Supernatant DNA amount for each concentration of Satb1 is plotted and shown. **D.** Droplet assays with constant protein concentration (0.5 μM) were incubated with the denoted concentrations of 1,6 hexanediol (1,6 HD) for 2 different time points and imaged.

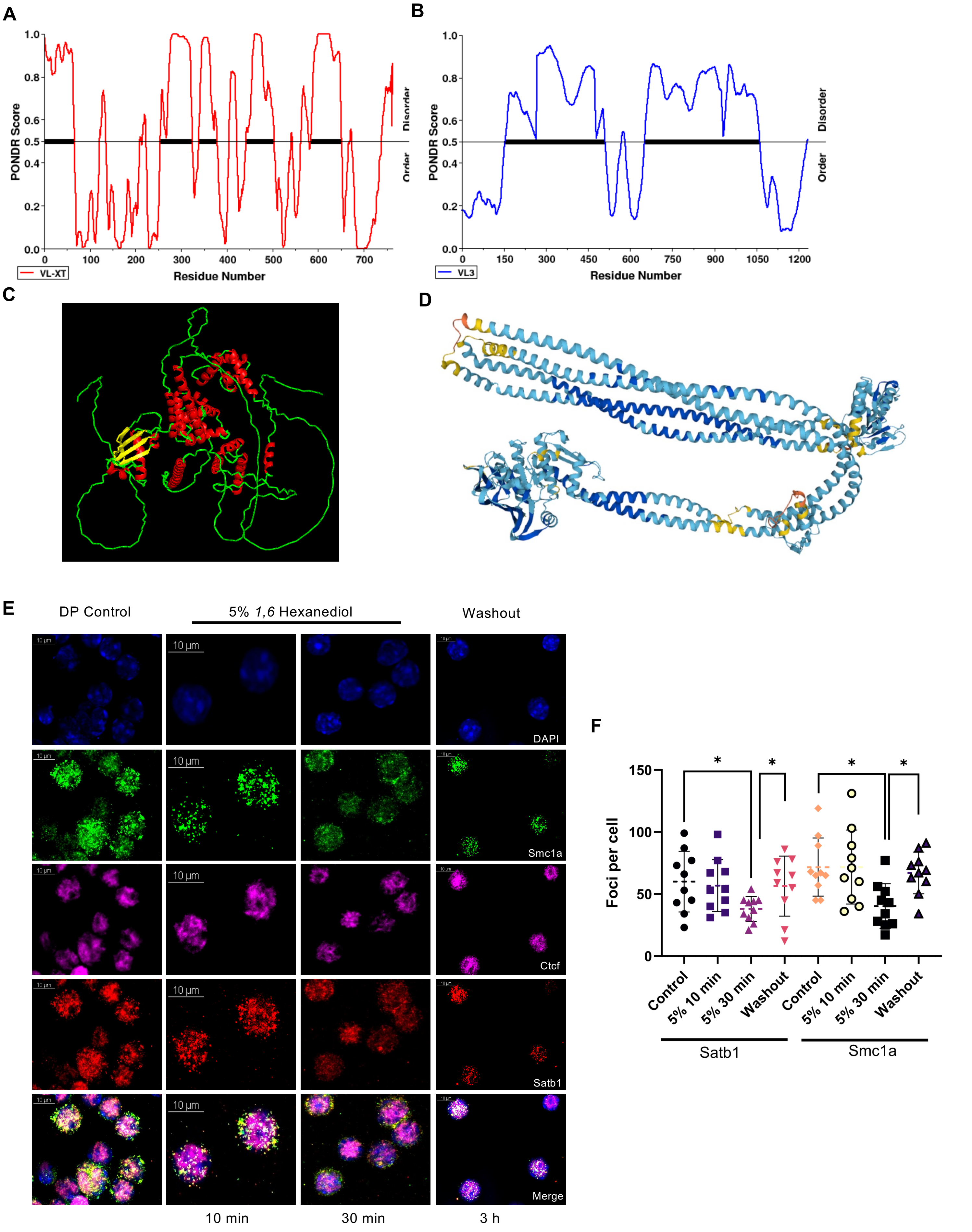

A
PONDR Score
Disorder
Order
Residue Number
VL-XT
B
PONDR Score
Disorder
Order
Residue Number
VL3
C
D
E
DP Control
5% 1,6 Hexanediol
Washout
10 µm
DAPI
Smc1a
Ctcf
Satb1
Merge
10 min
30 min
3 h
F
Foci per cell
Control
5% 10 min
5% 30 min
Washout
Satb1
Smc1a


Figure S13

**Figure S13. Satb1 condensates exhibit liquid-like property in DP thymocytes.** Disorder prediction of Satb1 (**A)** and Smc1a (**B**) amino acid sequences using PONDR webtool. **C** and **D.** Predicted Alphafold2 structures for Satb1 and Smc1a are shown as 3D models using PyMol. **E**. DP thymocytes were cultured in vitro with or without the presence of 1,6 HD. Immunostaining images shown for Satb1 (red), Smc1a (green) and Ctcf (magenta). **F.** Quantitation of the number of foci in Satb1 and Smc1a in control, 1,6 HD treated and 1,6 HD washed out samples. Representative images are shown for N=3. Graphpad v10 was used for statistical analysis, n=30. Two-sided ANOVA was performed * $p<0.05$, ns non-significant.

A

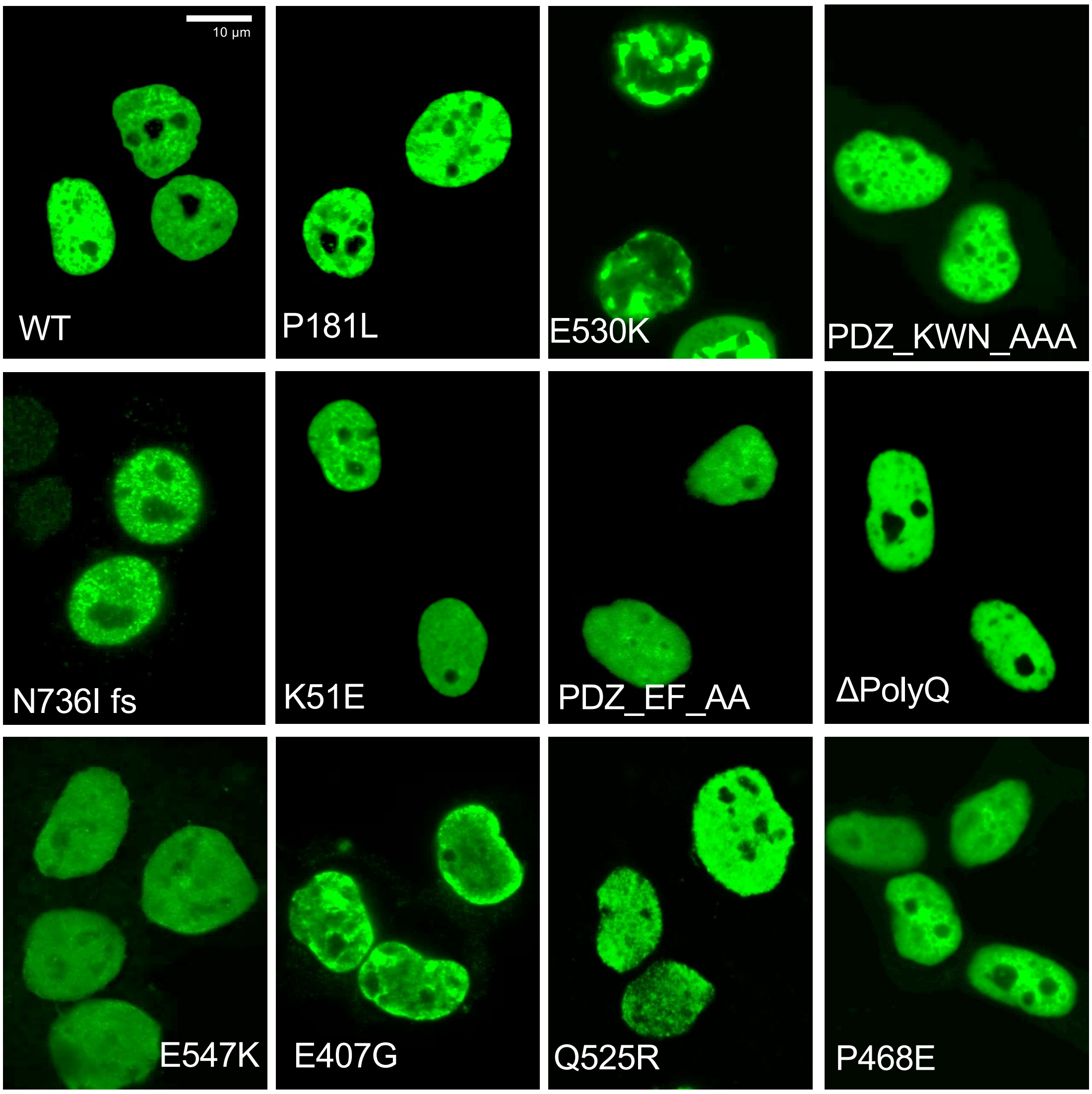


B

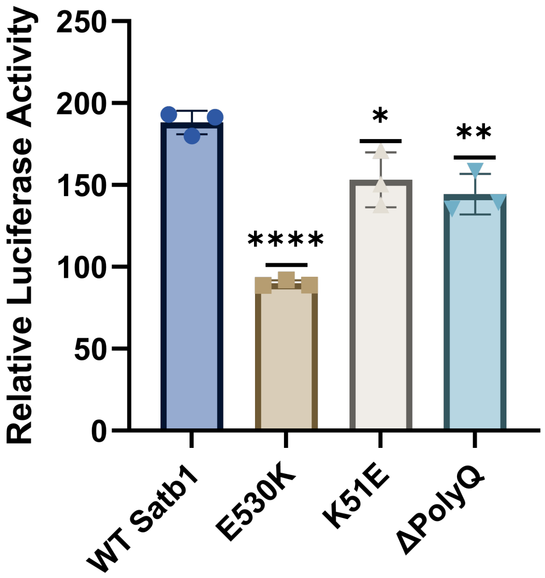


Figure S14

**Figure S14. Point mutations in SATB1 lead to dysregulation in its LLPS propensity. A.** Live-cell imaging of HEK-293T transfected with SATB1 GFP mutants and foci formed in each mutant are shown by confocal images. Representative of 3 independent transfections. **B.** Dual luciferase assay was performed upon overexpression of SATB1 mutants along with the reporter plasmid containing Il2 promoter. Luciferase was normalized to Renilla activity (pTRK) and shown for the most significantly affected mutants (indicated) compared to WT SATB1, n=4. Graphpad v10 was used for statistical analysis, two-sided ANOVA was performed * $p<0.05$, ** $p<0.01$, *** $p<0.001$.

| Gene (Forward and reverse) | Sequence |
|---|---|
| Cd8a F | CTTCCAGAACTCCAGCTCCA |
| Cd8a R | ACCGAGTTGCTGATGACTGA |
| Cd4b F | TTTGCAGAGGAAAACGGGTG |
| Cd4 R | AGAGTCAGAGTCAGGTTGCC |
| Cd3e F | CCTGAAAGCTCGAGTGTGTG |
| Cd3e R | TGGGCTCATAGTCTGGGTTG |

**Supplementary Table 1**: List of primers used for RT-qPCR (Related to Figures 5J, S9B).

| Primer | Sequences |
|---|---|
| Cd3e_promoter F | CTGAACCCCACACAGGAAGT |
| Cd3e_promoter R | TTTGCACAGGGCACAAGTAG |

**Supplementary Table 2**: List of *Cd3e* promoter-targeting primers used for ATAC-qPCRs (Related to Figure S10D).

| Mutation (Forward and reverse primers) | Sequence |
|---|---|
| P181L_F | CTTGCCTCTCGAACAATGGTCGCACAC |
| P181L_R | TGTTCGAGAGGCAAGTCTTCTAGTTTGGGG |
| E530K_F | GGTTGTGCAAGCTGTTACGCTGGAAAGAAGA |
| E530K_R | ACAGCTTGCACAACCATCCCTGGCT |
| E547K_F | CCCTGTGGAAGAACCTCTCCATGATCCGAAGGT |
| E547K_R | GGTTCTTCCACAGGGTTCTGTTTTCTGGAGA |
| E407G_F | GCTTTCAGGAATCCTCCGAAAGGAAGAGGACC |
| E407G_R | AGGATTCCTGAAAGCAAGCCCTGAGTTCTG |
| Q525R_F | CAAAAGCCGGGGATGGTTGTGCGAGCTG |
| Q525R_R | CATCCCCGGCTTTTGGTTGCTGCAAC |
| N736I fs_F | AGATAAAATACTAACACCCTTTTTTCAGTGAA |
| N736I fs_R | GTTAGTATTTTATCTTGGACACTCTCTTCCA |
| K51E_F | CAGGTGCAGAAATGCAGGGAGTGCCTTTAAAAC |
| K51E_R | GCATTTCTGCACCTGTACTCCCAAGCC |
| PDZ_EF_AA_F | CCAGCACAGCTGCTGCATGCTCCTCCTTGCAAT |
| PDZ_EF_AA_R | GCATGCAGCAGCTGTGCTGGTGAGAAAGGATAT |
| KWN136-138AAA FP | CTGGAGCCGCCGCTCCAACCTGGATTAGCCCT |
| KWN136-138AAA RP | GTTGGAGCGGCGGCTCCAGTTCCACTGTCTTACGT |
| DEL575-650 FP | GCTTCCACTGAAATCACCGCGTTGCTCTCCTGTTCAT |
| DEL575-650 RP | GCGGTGATTTCAGTGGAAGCCTTGGGAATCCTCCA |
| P468E FP | CAGCACACCAGAAAGCCGTCCTCCCCAGGTGAAAACA |
| P468E RP | GGACGGCTTTCTGGTGTGCTGATGAGGGGGGCAGGA |

**Supplementary Table 3:** List of oligonucleotide primers used for site-directed mutagenesis (related to Figures 7G-I, and S14).

**Supplementary Video 1. Proximity Ligation Assay (PLA) reveals interaction between Satb1 and Smc1a in thymocytes.** Yellow puncta represent amplified PLA signals marking sites of Satb1–Smc1a colocalization. Imaging was performed using a Nikon AXR NSPARC confocal microscope at 100X magnification with Z-stack image acquisition to obtain spatial resolution across the sample depth. Three-dimensional (3D) reconstruction and projection along the X, Y, and Z axes were subsequently performed using NIS-Elements AR software (v6.10) to visualize the spatial distribution of amplification signals within the sample. A still image from the movie is presented below.

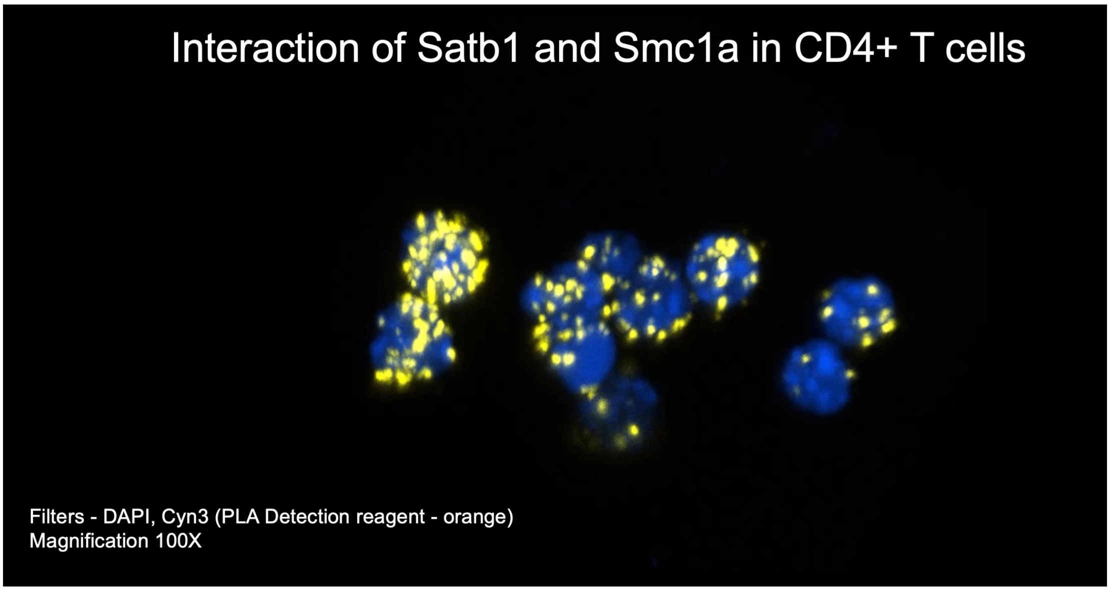
